# Modeling tissue-resident macrophage development from mouse pluripotent stem cells

**DOI:** 10.1101/2025.05.01.651743

**Authors:** Ann K. Baako, Ragavi Vijayakumar, Daniel Medina-Cano, Zhaoquan Wang, Jesús Romero-Pichardo, Kelvin Fadojutimi, Stephanie C. Do, Yuan Lin, Mohammed Islam, Sanjana Dixit, Alissa J. Trzeciak, Justin S.A. Perry, Thomas Vierbuchen

**Affiliations:** Developmental Biology Program, Sloan Kettering Institute, Memorial Sloan Kettering Cancer Center, New York, NY, USA; Immunology Program, Sloan Kettering Institute, Memorial Sloan Kettering Cancer Center, New York, NY, USA; Neuroscience Program, Weill Cornell Graduate School of Medical Sciences, Cornell University, New York, NY, USA

## Abstract

Tissue-resident macrophages (TRMs) are innate immune cells that participate in tissue development, homeostasis, and immune surveillance. Extensive efforts have been made to recapitulate TRM development from pluripotent stem cells (PSCs) *in vitro* to study molecular and cellular mechanisms of TRM development and to create cellular models of disease.

However, available PSC models of mouse TRM development exhibit low overall efficiencies of TRM generation, produce heterogeneous off-target populations, and rely upon undefined media components, thus limiting their reproducibility, scalability, and widespread application as an experimental platform for TRM biology. To address these important limitations, we developed an efficient and reproducible protocol to faithfully recapitulate the stepwise differentiation of mouse PSCs (epiblast stem cells) into unspecialized, proliferative TRMs through the pro-definitive hematopoietic program under defined conditions. These immature TRMs can stably integrate into developing mouse neural organoids *in vitro* and acquire features of microglia. In addition, PSC-derived immature TRMs can stably engraft into the lung niche *in vivo* and adopt alveolar macrophage characteristics. This new platform for modeling mouse TRM development represents a powerful experimental model system for studying TRM function and dysfunction in development and disease.

## INTRODUCTION

During early embryogenesis, tissue-resident macrophages (TRMs) – professional phagocytes of the innate immune system that are responsible for engulfing and digesting unwanted material – colonize nearly all developing organs. TRMs co-develop with their respective tissues, acquiring specialized functions that support the specific needs of each tissue during development and homeostasis ^1,2^. For example, in the brain, microglia regulate neural circuit formation by pruning excess synapses and clearing apoptotic cells ^3,4^. In the lung, alveolar macrophages clear inhaled material and recycle surfactant to maintain proper respiratory function throughout life ^5,6^. It is now appreciated that TRMs also play important roles in the pathophysiology of many diseases, including Alzheimer’s disease, schizophrenia, and atherosclerosis ^1,7^. Given the diverse and widespread functions of TRMs in tissues throughout the body and in various pathological states, there is significant interest in identifying strategies to modulate TRM function and to generate specific TRM populations from pluripotent stem cells (PSCs) for applications in regenerative medicine ^8–11^.

Pioneering studies in mouse and human models have established that TRMs originate from erythro-myeloid progenitors (EMPs) born in the yolk sac (referred to as yolk sac hematopoiesis or pro-definitive hematopoiesis) rather than from bona fide hematopoietic stem cell (HSCs), which are generated slightly later during definitive hematopoiesis ^1,12,13^. In mice, early mesodermal cells are generated from the posterior primitive streak region and migrate into the developing yolk sac around embryonic day 6.5-7.0 (E6.5-E7.0), where they differentiate into extra-embryonic mesoderm ^14,15^. Extra-embryonic mesoderm further differentiates into yolk sac mesoderm that gives rise to the yolk sac endothelium, a subset of which will begin to generate hemogenic endothelial cells beginning at ∼E8.25 ^2,16,17^. Yolk sac hemogenic endothelial cells undergo an endothelial-to-hematopoietic transition (EHT) to produce EMPs, which can either differentiate locally in the yolk sac into pre-macrophages (pMacs) or enter the nascent circulation (after E9.5) and colonize the fetal liver ^18,19^. Within the fetal liver, EMPs differentiate into pMacs as well as other hematopoietic progeny that support embryonic development, including erythrocytes, megakaryocytes, granulocytes, and mast cells ^1,17–23^. pMacs exit the fetal liver and migrate throughout the embryo, seeding developing tissues, undergoing local expansion, and then further differentiating into specialized, mature TRMs in response to tissue-specific instructive cues. Importantly, some TRM populations, such as microglia and Langerhans cells (TRMs that inhabit skin and other epithelial tissues), persist throughout life in mice with minimal replacement from infiltrating HSC-derived macrophages ^24–27^.

The complex and dynamic nature of yolk sac hematopoiesis makes it challenging to characterize the underlying molecular and cellular mechanisms. *In vitro* differentiation of pluripotent stem cells (PSCs) into yolk sac hematopoietic progenitors has emerged as a powerful experimental system for modeling TRM development ^28–31^. For example, human PSC (hPSC) protocols can recapitulate features of yolk sac hematopoiesis, generating pre-hematopoietic mesoderm, hemogenic endothelial cells, EMPs, immature TRMs, and specialized TRMs such as microglia-like cells ^32–37^. These reduced complexity models have helped to delineate critical signaling pathways and transcriptional regulatory mechanisms that orchestrate specification of mesodermal populations into yolk sac hemogenic endothelium and EMPs ^34,35,38–40^. In addition, generation of human TRMs *in vitro* can be used to generate cellular models of TRM function and disease biology and can serve as a potential source of cells for next-generation cell therapies ^33,35,41^.

Although much of our understanding of TRM development and function has come from studies in mouse models, protocols for differentiating mouse PSCs into TRMs remain significantly less advanced compared to analogous protocols for human PSCs, and thus they have not been widely adopted. For example, current protocols for modeling mouse yolk sac hematopoiesis generate heterogeneous populations of mesoderm in 3D embryoid bodies, require purification of intermediate populations to achieve a significant yield of desired populations, and utilize undefined media components such as serum ^42–44^. In contrast, there are now hPSC protocols for stepwise modeling of yolk sac hematopoiesis that more closely recapitulate the relevant developmental transitions, generate desired populations with moderate/high efficiency, and use defined media components ^35^.

The gap in protocol development between mouse and human PSC systems can be attributed in part to differences in PSC culture methods commonly used for each species. Mouse PSCs are typically cultured in the naïve pluripotent state, whereas human PSCs are cultured in the primed pluripotent state ^45–47^. As a result, naïve mouse PSCs must first transition from a naïve to a primed pluripotent state to effectively respond to differentiation signals and differentiate into germ layer progenitors ^46,48^. This naïve-to-primed transition involves rapid proliferation and is inherently asynchronous, thus introducing significant heterogeneity at this initial stage that complicates the subsequent stages of any differentiation protocol ^49^. Recent work from our group and others suggests using mouse primed-state PSCs (epiblast stem cells [EpiSCs]) as a starting point for directed differentiation can overcome some of these issues, enabling the development of robust and efficient differentiation models ^50–52^.

Here, we establish a defined protocol for the stepwise specification of mouse EpiSCs into EMPs and subsequently into a > 90% pure population of immature TRMs in 2D adherent culture without the need for purification or co-culture with other cell types. By optimizing the timing and strength of Wnt and TGF-β signaling, we identify conditions for efficient specification of pre-hematopoietic mesoderm (PHM), leading to increased yields of EMPs and immature TRMs.

This protocol is reproducible across multiple genetically diverse PSC lines and can be readily scaled to generate large quantities of EMPs and immature TRMs. Immature TRMs exhibit surface markers, gene expression profiles, and functional properties characteristic of yolk sac-derived TRMs *in vivo*. These immature TRMs can integrate into mouse cerebral cortex organoids and the *in vivo* lung niche, where they adopt features of mature TRMs. This new platform for modeling mouse TRM development will enable paired *in vitro* and *in vivo* studies that leverage the full suite of mouse genetic tools, and thus should be a powerful experimental model system for mechanistic studies of TRM function and dysfunction in development and disease.

## DESIGN

Current state-of-the-art protocols for modeling yolk sac hematopoiesis and generating TRMs from mouse PSCs have significant shortcomings compared to analogous protocols that have been developed for human PSCs. First, current protocols start from PSCs cultured in the naïve pluripotent state (mouse embryonic stem cells), which must first transition into a primed pluripotent state before further differentiation can proceed ^45,46^. This initial step is inherently stochastic, making downstream differentiation stages difficult to control. Additionally, cells at this stage exhibit high proliferation rates and increased frequency of apoptosis, further complicating differentiation efforts ^46^. Second, current protocols perform differentiation in 3D embryoid body (EB) cultures, which are more heterogeneous compared to 2D adherent cultures. This heterogeneity impacts reproducibility, leading to variable differentiation outcomes across technical replicates and different cell lines. The heterogeneity and asynchronous nature of differentiations also complicates efforts to model intermediate steps of pro-definitive hematopoiesis. Finally, many existing protocols rely on serum, which contains complex, undefined components that can introduce batch-to-batch variability, making it more difficult to study the specific signaling pathways and other cues critical for each stage of differentiation. Given these shortcomings, we sought to develop an improved protocol for modeling mouse yolk sac hematopoiesis (pro-definitive hematopoiesis) and TRM differentiation *in vitro*. Such a model would be of interest to researchers studying TRM biology and development *in vivo* as it would serve as a complementary reductionist model for in-depth mechanistic studies.

To establish a new mouse model of yolk sac hematopoietic differentiation, we began by adapting a recently developed 2D, serum-free protocol for generating yolk sac-derived TRMs from hPSCs ^35^. hPSCs are generally cultured in the primed pluripotent state, corresponding to the pluripotent cells in the post-implantation epiblast, which are competent to directly differentiated into progenitors of each of the three embryonic germ layers ^45,53^. Mouse PSCs can also be cultured in the primed pluripotent state (EpiSCs), but mouse EpiSCs have not been broadly adopted as a starting point for *in vitro* directed differentiation studies ^47,54^. We recently demonstrated that hPSC protocols can be adapted for the differentiation of mouse EpiSCs into definitive endoderm and neuroectoderm ^50,55^. Based on these results, we chose to adapt an hPSC TRM differentiation protocol for directed differentiation of mouse EpiSCs into TRMs ^35^.

This required careful consideration of species-specific differences, including 1) mouse development progresses more rapidly than human development, 2) extra-embryonic mesodermal progenitors experience distinctive signaling environments in mouse compared to human due to differences in yolk sac development, and 3) the surface marker profile used to define early yolk sac mesoderm subpopulations in human does not translate directly to mouse. Consequently, adapting the protocol involved refining timing and developmental cues and validating markers to align with mouse-specific developmental processes.

## RESULTS

### Identification of conditions for generating yolk sac pre-hematopoietic mesoderm (PHM) from mouse epiblast stem cells

Starting at ∼E6.25, pluripotent cells in the epiblast ingress through the posterior primitive streak and give rise to nascent mesoderm cells, a subset of which migrate proximally towards the extra-embryonic ectoderm to become extra-embryonic mesoderm ^14,15,56^. Extra-embryonic mesodermal cells further differentiate into several distinct populations by E7.75-E8.0, including the allantois core domain, allantois outer mesenchyme, and yolk sac mesoderm ^57^. The yolk sac mesoderm can be distinguished by expression of FOXF1 and GATA6, and by the absence of T/BRA and SOX17 ^57^. A subset of yolk sac mesoderm further differentiates into arterial and venous endothelial cells that make up the yolk sac endothelium, and some of these endothelial cells further differentiate into hemogenic endothelium ^58–60^. Finally, hemogenic endothelial cells undergo an endothelial-to-hematopoietic transition (EHT) to produce multipotent erythro-myeloid progenitors (EMPs), which give rise to embryonic erythroid and myeloid lineages, including tissue-resident macrophages ^61^ (**Fig. 1A**). We sought to recapitulate these stepwise developmental transitions, starting from mouse epiblast stem cells, which are equivalent to pluripotent epiblast cells in the pre- or peri-gastrulation mouse embryo (**Fig. 1B, Supplementary Fig. 1A**).

**Figure 1:**
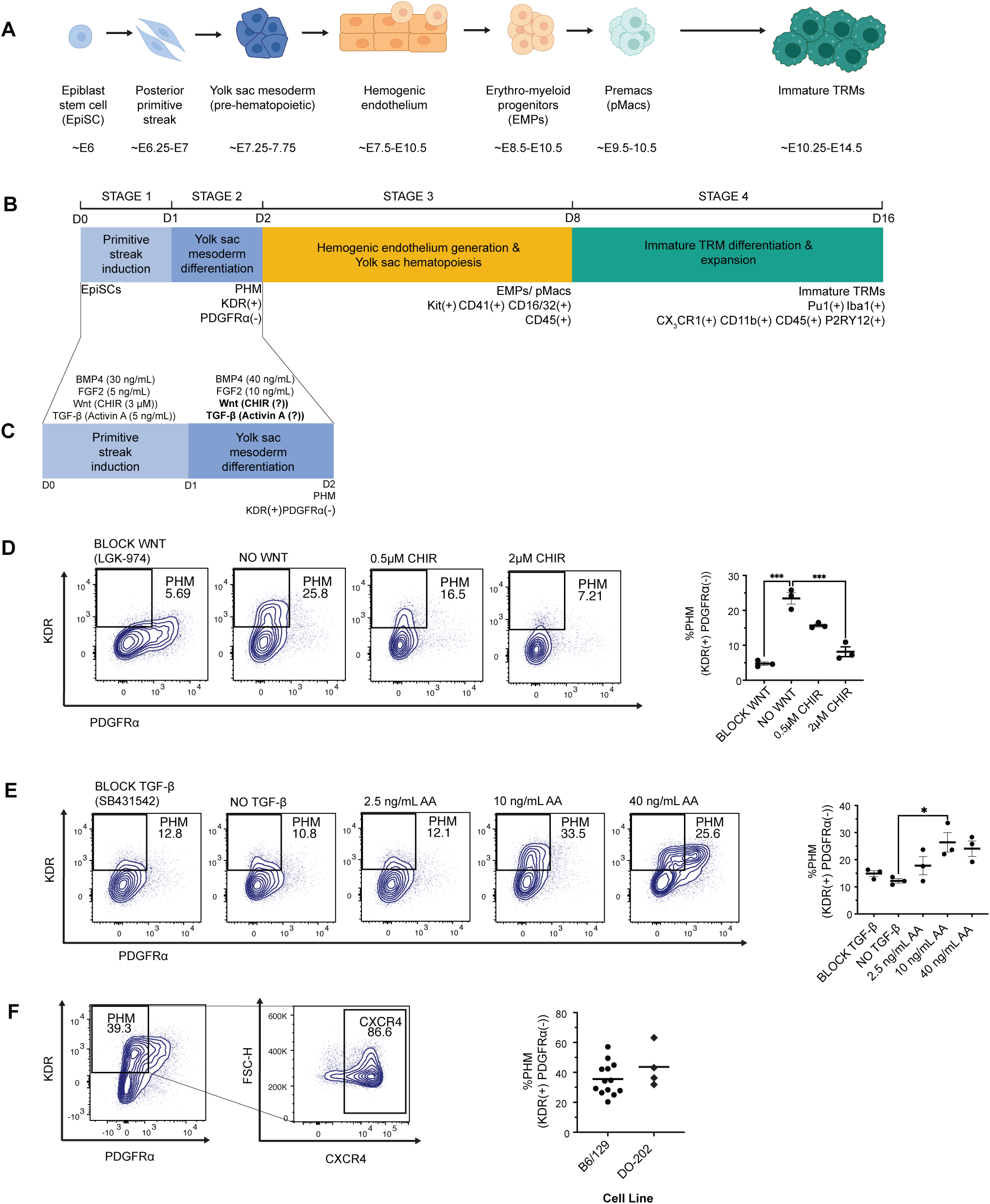
Identification of conditions for generating yolk sac pre-hematopoietic mesoderm (PHM) from mouse EpiSCs. **A**. Overview of developmental stages from pluripotency through yolk sac hematopoiesis and TRM differentiation, including relevant stages of embryonic development in mouse (bottom row). **B**. Overview of approach for differentiation of EpiSCs into immature TRMs, including stage-specific marker genes. **C**. Summary of the first two stages of the TRM differentiation protocol. The optimal levels of Wnt and TGF-β signaling to promote KDR(+) PDGFRa(-) yolk sac mesoderm/PHM differentiation during Stage 2 were not known. **D**. Representative flow cytometry plots (left) of D2 PHM yield observed at different levels of Wnt pathway activation. CHIR99021 was used to activate the canonical Wnt pathway, and LGK479 was used to inhibit Wnt signaling. Activin A (AA) was kept constant at 10 ng/mL during stage 2 for each condition. Summary of flow cytometry data across 3 independent differentiation experiments (right). Significance was determined using ordinary one-way ANOVA, corrected for multiple comparisons with Tukey test (***P < 0.001). **E**. Representative flow cytometry plots (left) showing D2 PHM yield observed at different levels of TGF-β pathway activation. Activin A was used to stimulate TGF-β signaling and SB431542 was used to block TGF-β signaling. Only endogenous Wnt signaling (same as the No Wnt condition in [D] was used in Stage 2). Summary of flow cytometry data across 3 independent differentiation experiments (right). Significance was determined using ordinary one-way ANOVA, corrected for multiple comparisons with Tukey test (*P < 0.05; **P < 0.01; ***P < 0.001). **F**. Representative flow cytometry plots (left) measuring PHM yield at the end of Stage 2 in optimized PHM conditions (No Wnt, 10 ng/mL Activin A). Percentage of CXCR4 within the KDR(+) PDGFRa(-) population is also shown. Summary of flow cytometry data from 2 distinct cell lines (right). B6/129 EpiSCs (n = 13 independent differentiations [technical replicates]). DO-202 EpiSCs (n = 4 independent differentiations [technical replicates])

However, in mice, the exact surface marker profile of yolk sac PHM that gives rise to hemogenic endothelium remains incompletely defined, complicating the assessment of specification efficiency during early differentiation stages. Previous studies in mouse embryos and in embryoid bodies derived from mouse PSCs found that the KDR^+^ PDGFRα⁻ population is enriched for cells with hematopoietic potential ^42,44,62–64^. Furthermore, a recent study reported that mouse and human PHM cells are KDR^+^ CXCR4^+^ ^65^. Based on these data, we chose to use KDR^+^ PDGFRα⁻ and KDR^+^ CXCR4^+^ marker profiles for evaluating the efficiency of PHM induction from EpiSCs.

We first sought to identify the appropriate signaling conditions to specify PHM from EpiSCs. We began by differentiating EpiSCs into T/Bra^+^ posterior primitive streak cells using an established serum-free cocktail of growth factors – FGF2, TGF-β (Activin A), BMP4, and Wnt (CHIR99021) ^35,36,66^. This generates an average of ∼80% T/Bra^+^ posterior primitive streak-like cells in 24 hours (n = 3 distinct EpiSC lines) (**Fig. 1C and Supplementary Fig. 1A-C**). To further specify posterior primitive streak progenitors into KDR^+^ PDGFRα⁻ PHM, we maintained high levels of BMP4, given its expression throughout the extra-embryonic ectoderm during gastrulation ^14^, and examined the impact of modulating the levels of Wnt and TGF-β signaling during the second day of differentiation.

We first modulated Wnt signaling during day 2 (D2) and evaluated KDR^+^ PDGFRα⁻ PHM generation. To do this, we kept Activin A constant at 10 ng/mL and FGF2 and BMP4 at 10 ng/mL and 40 ng/mL respectively. We tested four different Wnt conditions – Block Wnt (4 nM LGK-974), endogenous Wnt signaling only (no addition of Wnt inhibitor and no addition of CHIR99021), moderate Wnt activation (0.5 μM CHIR99021), and high Wnt activation (2 μM CHIR99021). We found that endogenous Wnt levels yielded the highest number of KDR^+^ PDGFRα⁻ cells (25.8%, **Fig.1C, D**). Interestingly, this result is distinct from what has been observed in hPSC differentiation, where it is critical to inhibit Wnt signaling shortly after primitive streak differentiation to specify PHM ^35,39,40^.

Having established the optimal level of Wnt signaling, we next investigated the effect of different levels of TGF-β pathway activity on the KDR^+^ PDGFRα⁻ population. These experiments indicated that 10 ng/mL of Activin A generates the highest frequency of KDR^+^ PDGFRα⁻ PHM cells, and that ∼87% of the KDR^+^ PDGFRα⁻ PHM cells in this condition also express CXCR4 (**Fig. 1C, E, F**). Further increasing Activin A (40 ng/mL) caused an increase in the KDR^+^ PDGFRα^+^ population, which is thought to be enriched for cardiogenic mesoderm ^42^. Interestingly, the frequency of KDR^+^ CXCR4^+^ cells is more sensitive to the levels of TGF-β signaling, with the highest fraction of KDR^+^ CXCR4^+^ cells observed at the highest levels of Activin A (40 ng/mL), and much lower levels observed in conditions where TGF-β is low or actively blocked (**Supplementary Fig. 1D**). Based on these results, we chose to proceed using the 10 ng/mL Activin A condition to maximize the KDR^+^ PDGFRα⁻ CXCR4^+^ population.

These optimized conditions generated ∼40% KDR^+^ PDGFRα⁻ PHM cells at 48 hours. Importantly, specification of PHM from EpiSCs was reproducible between technical replicates (the same cell line differentiated in independent experiments) and from a second EpiSC line from an outbred genetic background, DO-202 (B6/129 n = 13 technical replicates, DO-202 n = 4 technical replicates) (**Fig. 1F**). Further characterization of differentiating cultures at 48 hours revealed cells expressing markers of yolk sac mesoderm/PHM (FOXF1, GATA6) surrounding clusters of T/Brachyury^+^ cells, which are likely primitive streak-like cells or off-target extra-embryonic mesoderm subtypes that express T/Brachyury (**Supplementary Fig. 1E**). These optimized conditions for specification of PHM-enriched extra-embryonic mesoderm from EpiSCs provide a strong foundation for modeling subsequent stages of yolk sac hematopoiesis.

### Efficient generation of erythro-myeloid progenitors from pre-hematopoietic yolk-sac mesoderm

After optimizing conditions for generation of yolk sac PHM, we next sought to determine whether PHM can be further differentiated into yolk sac hemogenic endothelium and EMPs. To generate EMPs, we adapted differentiation conditions previously used for hPSC differentiation into EMPs, reducing the total time allotted for each step of differentiation to account for the faster pace of murine development compared to human. (**Fig. 2A**). We first induced hemogenic endothelium generation by adding VEGF to PHM cultures. Once hemogenic endothelium had been generated, we sequentially supplemented media with SCF, IL-6, TPO, and IL-3, cytokines known to induce hematopoiesis and promote EMP expansion ^35^. In the developing mouse embryo, EMPs emerge from yolk sac hemogenic endothelial cells via an endothelial-to-hematopoietic transition and can be isolated based on their expression of the surface markers Kit, CD41, and CD16/32 ^67^. Using this surface marker profile, we monitored the generation of EMPs in differentiating cultures between D2 and D7 (**Fig. 2B**). Semi-adherent EMPs were first observed at low frequency at D5 (∼6%) and peaked at ∼20% by D7-D8, comparable to the ∼17% EMP frequency reported in a similar hPSC-based model ^35^ . Under these conditions, essentially all EMPs also expressed the pan-myeloid marker CD45 and the transcription factor RUNX1, which is required for EMP generation from yolk sac hemogenic endothelium (**Fig. 2B, C**) ^68^. Importantly, we found that EMP generation was reproducible across technical replicates and in a second EpiSC line, DO-202, albeit at slightly lower efficiency (n = 4 independent differentiation experiments for each cell line, **Fig. 2B**). Taken together, these data demonstrate that PHM generated from EpiSCs can be efficiently and reproducibly differentiated into EMPs over a period of 3-5 days.

**Figure 2:**
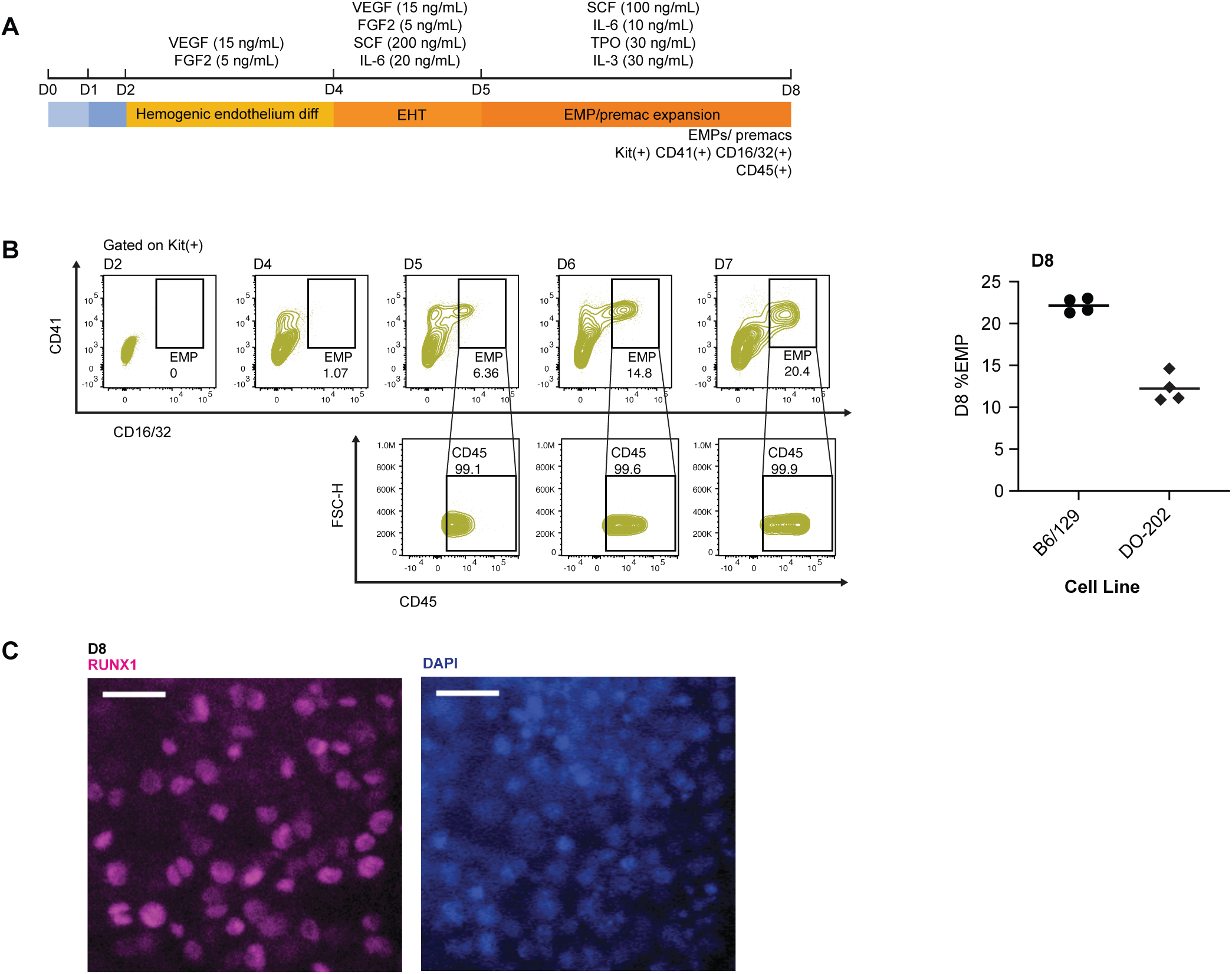
Efficient generation of erythro-myeloid progenitors from pre-hematopoietic yolk sac mesoderm. **A**. Detailed overview of signaling conditions for hemogenic endothelium differentiation and hematopoiesis. These conditions are adapted (with modifications) from Guttikonda et al., 2021. **B**. Evaluation of EMP emergence during in vitro hematopoiesis (left). Generation of Kit(+) CD41(+) CD16/32(+) EMPs and myeloid marker expression (CD45) was evaluated using flow cytometry at the indicated timepoints. Flow cytometry plots are gated for Kit(+) cells (top row). Under these conditions, essentially all EMPs also express CD45 (bottom row). Quantification of EMP generation (D8) across 4 independent differentiation experiments from the indicated cell lines (right). **C**. Immunofluorescence staining of differentiating cultures at D8 for RUNX1, a transcription factor expressed in yolk sac hemogenic endothelia and EMPs. EHT = endothelial-to-hematopoietic transition. Scale bars = 25 µM.

### Differentiation of EMPs into immature TRMs and characterization of immature TRMs

To further differentiate EMPs into pMacs and immature TRMs, we dissociated differentiating cultures at D8 and replated cells onto Matrigel-coated tissue culture plates in serum-free media containing M-CSF and IL-34, growth factors that both signal through the CSF1 receptor (CSF1R) to stimulate TRM differentiation and proliferation during embryonic development (**Fig. 3A**). After two days in these conditions (D10 overall), round, vacuolated, immature TRMs with filopodia were present throughout the culture (**Fig. 3B**). Immature TRMs are highly proliferative and continue to divide for at least 8 days under these conditions, giving rise to nearly pure populations of TRMs that express the pan-macrophage markers PU.1 and IBA1 by D16 (**Fig. 3B**). In addition, D16 TRMs uniformly express the pan-myeloid marker CD45 and the pan-macrophage markers CD11b and CX_3_CR1. A subset of TRMs (∼16%) express P2RY12, a receptor that is preferentially expressed on microglia and on some Kupffer cells *in vivo* ^69–71^ but generally not highly expressed on TRMs *in vitro* ^72–74^. Importantly, we observed efficient generation of CX_3_CR1^+^ cells at D16 across multiple, independent differentiations from three distinct EpiSC lines (B6/129 [n = 6 technical replicates from independent differentiations], DO-202 [n = 3 technical replicates from independent differentiations], and DO-223 [n = 3 technical replicates from independent differentiations], **Fig. 3C**). Beginning with 20,000 EpiSCs in a 96-well plate, each differentiation yielded approximately 360,000 CX_3_CR1^+^ immature TRMs per cell line.

**Figure 3:**
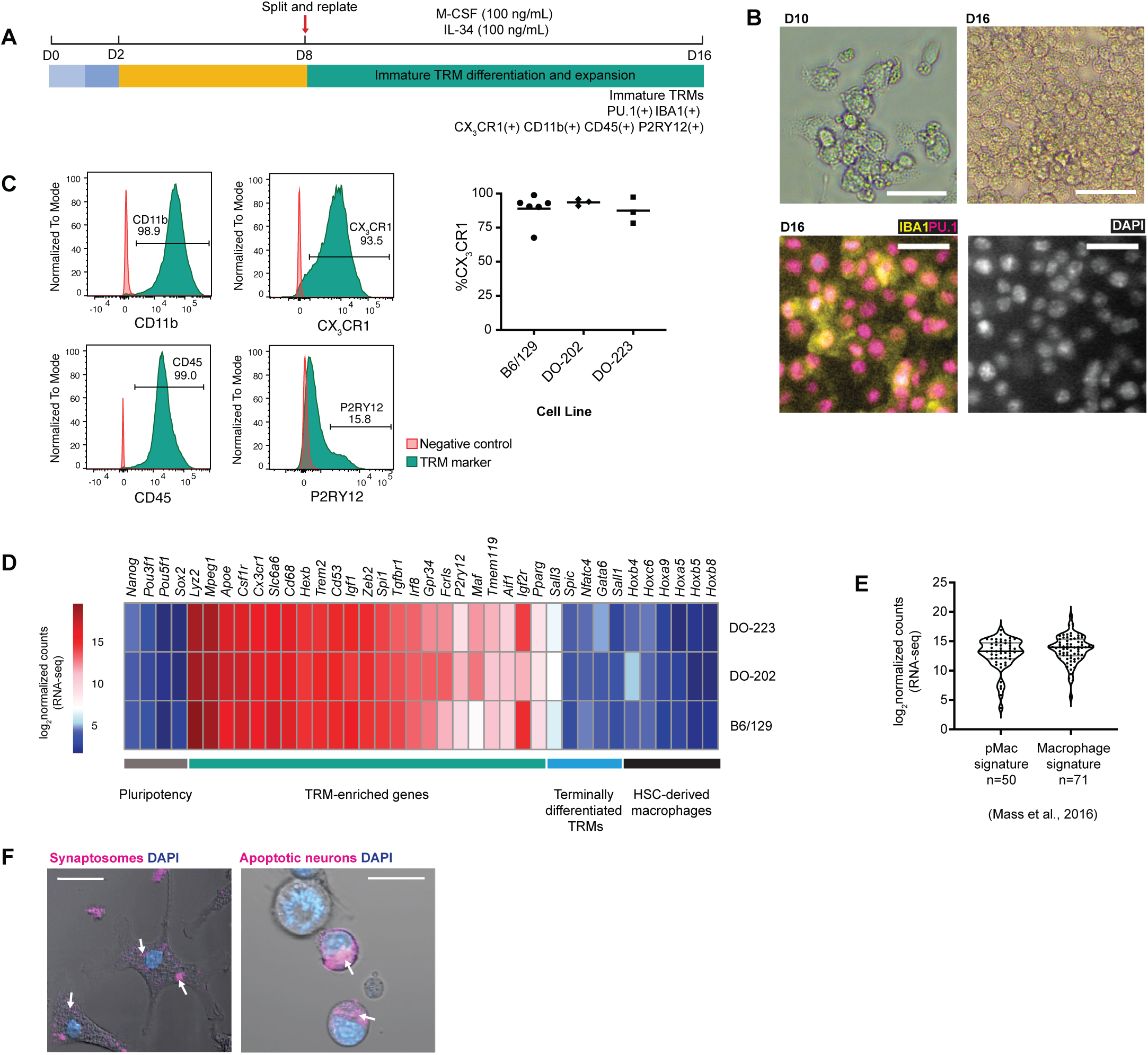
Differentiation of EMPs into immature TRMs and characterization of immature TRMs. **A**. Overview of conditions used for differentiation of EMPs into immature TRMs and TRM expansion. **B**. Brightfield images of immature TRMs at D10 and D16 (top row) and immunofluoresence staining immature TRMs (D16) for pan-macrophage markers IBA1 and PU.1 (bottom row). **C**. Assessment of TRM surface markers CX_3_CR1, CD11b, CD45, and P2RY12 by flow cytometry to evaluate immature TRM differentiation efficiency (left). Summary of differentiation efficiency (% CX_3_CR1(+) cells) obtained from the indicated EpiSC lines (right). Each dot represents an independent differentiation experiment. **D**. Heatmap of gene expression (RNA-seq) of a panel of cell type-specific marker genes in CX_3_CR1(+) immature TRMs purified by FACS at D16. Log2 normalized counts (transcripts per million [TPM]) are plotted for each gene (average expression level from n = 2 independent differentiations [technical replicates] from each of the indicated cell lines). **E**. Violin plots of expression levels (log2 normalized counts, TPM) of individual genes from signatures of yolk sac-derived pMacs and yolk sac-derived macrophages obtained from E10.25 and E10.5 embryos (Mass et. al., 2016). Each dot represents a single gene, and the value for the expression level of the gene is the average expression across TRMs from all of the 3 EpiSC lines. **F**. Confocal microscopy images (brightfield overlaid with fluorescence) of D20 immature TRMs containing pH-sensitive CypHer 5E-labeled phagocytosed synaptosomes (left) and apoptotic neurons (right). Arrows indicate engulfed cellular material. Scale bars = 25 µm.

To further evaluate the properties of immature TRMs, we performed RNA-seq on CX_3_CR1^+^-sorted immature TRMs from three distinct EpiSC lines (B6/129, DO-202, DO-223; n = 2 technical replicates per cell line). Consistent with the flow cytometry data, immature TRMs exhibited robust expression of macrophage-enriched genes (*Csf1r*, *Mpeg1*, *Slc6a6*, *Trem2*) as well as modest expression of TRM-enriched genes including *Tmem119* and *P2ry12* (**Fig. 3D**). In contrast, genes that are expressed specifically in mature, functionally specialized subtypes of TRMs, such as *Sall1* (microglia), *Nfatc4* (kidney TRMs), and *Spi-c* (spleen TRMs) were expressed at negligible levels, consistent with prior work indicating that TRMs *in vitro* fail to express markers characteristic of mature TRMs *in vivo* ^72–74^ (**Fig. 3D**). Immature TRMs also do not express Hox genes (*Hoxa5, Hoxa9, Hoxb4, Hoxb5, Hoxb8, Hoxc6*) that are normally expressed by HSC-derived macrophages ^75–77^, consistent with their generation via EMPs in the pro-definitive hematopoietic program (**Fig. 3D**). In addition, to evaluate whether immature TRMs were more similar to pMacs or embryonic TRMs, we examined the expression of gene signatures characteristic of *in vivo* pMacs from E9.5 and E10.25 yolk sac and head (n = 50 genes) and macrophages from E10.25 and E10.5 yolk sac and head (n = 71 genes, **Fig. 3E**) ^19^. *In vitro*-derived immature TRMs shared transcriptional features with both *in vivo* pMacs and embryonic macrophages, supporting their identity as an intermediate TRM state fated to mature into specialized TRMs.

We then examined the functional capacity of immature TRMs to phagocytose CypHer 5E-labeled substrates. The immature TRMs phagocytosed synaptosomes and apoptotic neurons within 45 minutes of exposure to the substrates (**Fig. 3F, Supplementary Video**).

To evaluate the potential of immature TRMs to acquire features of more specialized, tissue-specific macrophage populations *in vitro,* we chose to culture immature TRMs in conditions that promote acquisition of some features of microglia identity. To do this, at D11 we switched immature TRMs into media containing IL-34, TGF-β, and cholesterol based on a serum-free cocktail (TIC media) that was developed to promote primary microglia survival *in vitro* ^72^.

Specifically, we made a slight modification to this media composition by maintaining our macrophage base media but supplementing with the same concentrations of the key cytokines, lipids and acids found in the original TIC media. We refer to our media as the B27-TIC media and draw comparisons with two other media compositions – the original TIC media and immature TRM media (**Supplementary Fig. 2A**).

Both TIC and B27-TIC media conditions led to reduced proliferation and enhanced ramification of TRMs, whereas cells in immature TRM conditions remained rounded and continued to proliferate (**Supplementary Fig. 2A**). The highest degree of ramification was observed when cells in the two TIC conditions were split at D16 and replated onto collagen-coated tissue culture plates (**Supplementary Fig. 2A**). Surprisingly, we observed that attachment and survival of TRMs/microglia-like cells was significantly better in the B27-TIC media compared to the TIC condition, and thus we chose to use B27-TIC for subsequent experiments. We then compared transcriptomes of immature TRMs and microglia-like cells from three distinct EpiSC lines by isolating CX_3_CR1^+^ cells by FACS in each condition and performing bulk RNA-seq. Principal component analysis (PCA) revealed that TRMs in each condition separated by cell line on PC1, and then by media condition on PC2, suggesting that the differences in transcriptome between genetically distinct EpiSC lines were larger than the differences between TRMs in each of the two media conditions (**Supplementary Fig. 2B**). Compared to immature TRMs, the B27-TIC microglia-like cells expressed higher levels of *in vivo* microglia signature genes, including *P2ry12*, *Tmem119*, *Mertk*, *Tgfbr1*, *Hexb*, and *Sall3* (**Supplementary Fig. 2C**). B27-TIC microglia-like cells also had strong expression of genes expressed in primary microglia isolated from the brain at E14.5 and postnatal day 4/5 (P4/5) ^78^, further suggesting that they can acquire features of immature microglia (**Supplementary Fig. 2D**).

To compare B27-TIC microglia-like cells to primary microglia more directly, we isolated primary CD11b^+^ microglia from P4 mice and cultured them in B27-TIC for 6 days. We then examined expression of surface markers broadly expressed by TRMs (CD45, CD11b, CX_3_CR1) and more specifically enriched on microglia (P2RY12). Cultured primary microglia and B27-TIC microglia-like cells expressed similar levels of CX_3_CR1, CD11b, and P2RY12 within their respective CD45^+^ populations (**Supplementary Fig. 2E**). In addition to their transcriptional and phenotypic similarities to neonatal microglia, microglia-like cells displayed functional properties similar to their *in vivo* counterparts. Microglia-like cells successfully phagocytosed synaptosomes, apoptotic Jurkat cells, and the yeast antigen zymosan. Fixed confocal imaging confirmed that the microglia-like cells also expressed CD68, a marker of macrophage activation (**Supplementary Fig. 2F**). Thus, by culturing immature TRMs in conditions that more closely mimic the CNS environment, immature TRMs generated from mouse EpiSCs can be further induced to acquire some features of microglia, exhibiting general similarity to *ex vivo* cultured microglia from neonatal mice.

### Immature TRMs integrate into developing cortical organoids and adult mouse lung niche

The terminal differentiation and maintenance of functionally specialized TRMs requires continuous instructive signaling from their unique tissue microenvironments. Previous studies suggest that these tissue-specific signals are difficult to replicate in simple tissue culture models, and thus *ex vivo* and PSC-derived TRMs are more similar to immature, unspecialized TRMs ^72^. More complex models of developing tissues, such as 3D tissue organoids, can more closely recapitulate the complex signaling environment found in tissues *in vivo*. Therefore, we next sought to determine if immature TRMs can colonize mouse cerebral cortical organoids and acquire characteristics of microglia. We used a cerebral cortical organoid differentiation protocol established in our laboratory to generate cortical organoids ^55^. These organoids contain only neuroectodermal cell types, and thus do not contain microglia (**Supplementary Fig. 3**). We added immature TRMs (D28) to D10 cortical organoids, a time point that corresponds approximately to the mid-to-late stages of cortical neurogenesis *in vivo* (∼E15.5-E17.5). Cortical organoids with integrated TRMs were then cultured for an additional 14-21 days in cortical organoid media supplemented with IL-34 and TGF-B to promote proliferation and microglial differentiation of transplanted immature TRMs (**Fig. 4A**). Two weeks post-transplantation, approximately 7% of the cells within the cortical organoids expressed the microglia marker CD11b, with this population increasing to around 40% after an additional week of culture (**Fig. 4B**). The relative increase in abundance of transplanted TRMs over time likely reflects both microglial proliferation and death of neuronal and glial cells within the organoids. Immunofluorescence staining of tissue-cleared cortical organoids demonstrated that transplanted TRMs successfully integrated into the neuroepithelium of the developing organoids, proliferated, migrated throughout the organoids to tile the developing neuroepithelial tissue, and acquired ramified morphologies similar to microglia in developing cerebral cortex *in vivo* (**Fig. 4C, Supplementary Fig. 3**). These experiments provide a proof-of-concept that mouse EpiSCs can be used to generate chimeric cerebral cortical organoids containing microglia-like TRMs.

**Figure 4:**
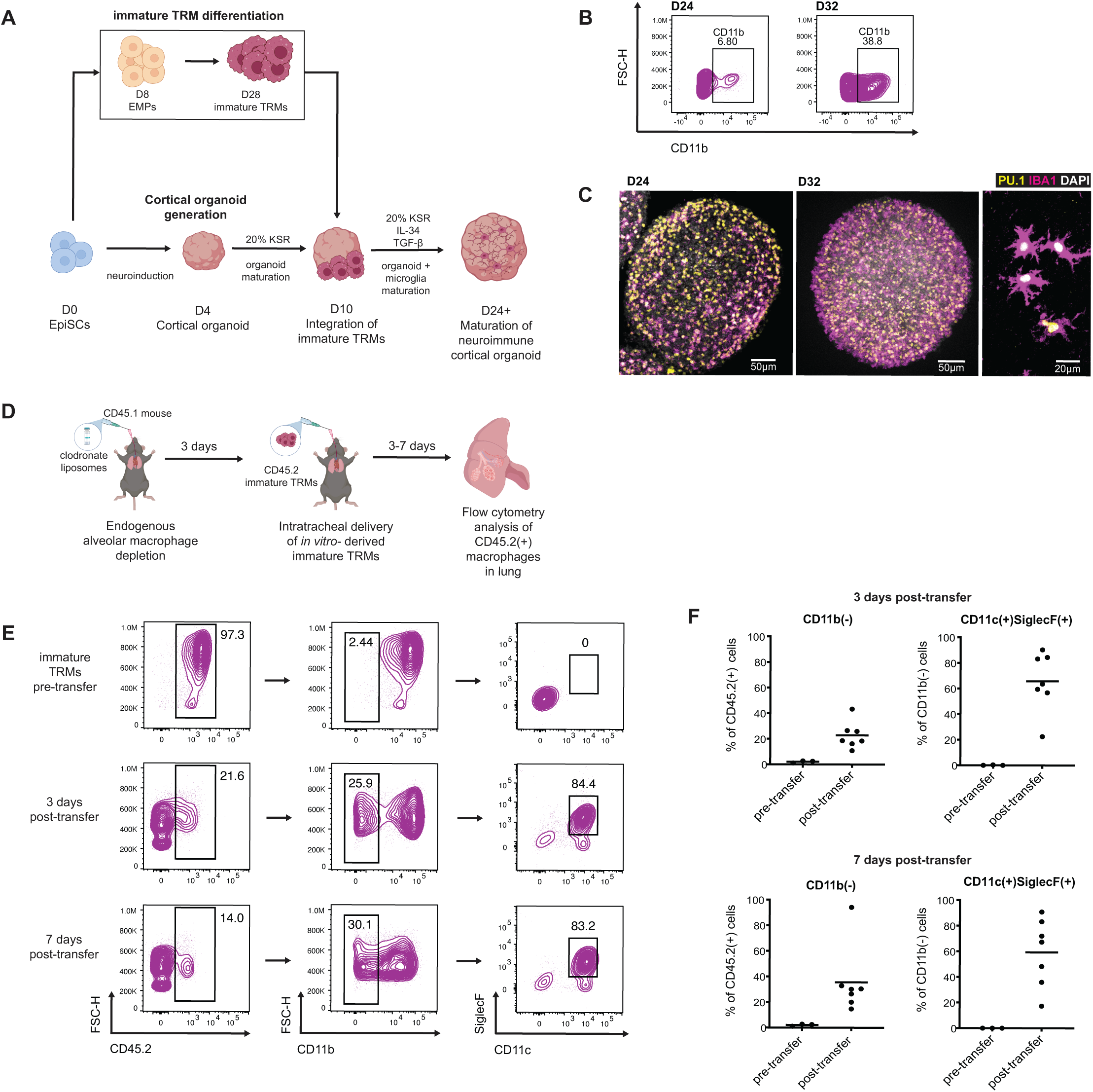
Immature TRMs integrate into developing cortical organoids and adult mouse lung niche. **A**. Overview of strategy for co-culture of immature TRMs and mouse cerebral cortex organoids generated from EpiSCs. **B.** Quantification of CD11b(+) TRMs in cortical organoids using flow cytometry at D24 and D32. **C**. Confocal imaging of tissue-cleared cortical organoids with integrated TRMs. Immunofluorescence staining for TRM markers PU.1 and IBA1. High magnification image (right) showing ramified morphology of TRMs. **D**. Overview of experimental design for depletion of endogenous alveolar macrophages and subsequent intratracheal delivery of immature TRMs into mouse lungs. CD45.1 mice were used so that transplanted CD45.2-expressing TRMs can be distinguished from CD45.1(+) host macrophages. **E**. Representative flow cytometry plots for lung tissue from mice transplanted with TRMs at the indicated timepoints. Alveolar macrophages are CD11b(-) SiglecF(+) CD11c(+). **F**. Summary of results from lung transplantation experiments at D3 and D7 post-transplant. Individual dots represent results from a single mouse (biological replicates).

Finally, we sought to assess the capacity of immature TRMs to engraft into specific tissue niches in adult mice. We selected the lung as our model due to the availability of a well-established, efficient protocol for adoptive cell transfer. We first depleted the endogenous alveolar macrophage population in CD45.1 congenic mice using clodronate liposomes ^79^. We then transplanted immature CD45.2^+^ TRMs intratracheally and allowed them to colonize lung tissue for 3 days or 7 days (**Fig. 4D**). By day 3, approximately 25% of the exogenous CD45.2 TRMs present in the lungs had reduced CD11b expression, which is typically observed in alveolar macrophages. Among these CD45.2^+^ CD11b⁻ cells, approximately 65% also expressed the alveolar macrophage surface markers CD11c and SiglecF, suggesting that transplanted immature TRMs had successfully integrated into the alveolar niche and acquired features of differentiated alveolar macrophages. Similar results were observed 7 days after transplantation, suggesting stable engraftment of EpiSC-derived TRMs (**Fig. 4E, F**).

Taken together, these data demonstrate that immature TRMs can integrate into two distinct specialized tissue environments and acquire characteristics of functionally specialized, mature TRMs, highlighting the potential utility of PSC-derived immature TRMs for studying TRM development and function in various tissue contexts.

## DISCUSSION

Here, we developed a defined (serum-free), adherent (2D) protocol for recapitulating mouse yolk sac hematopoiesis (pro-definitive hematopoiesis) starting from primed pluripotent stem cells (EpiSCs). This protocol can generate > 90% pure population of immature TRMs in 16 days without the need for purification or co-culture with other cell types. This new protocol addresses several of the limitations of EB-based approaches, leading to improved differentiation efficiency, scalability, and reproducibility across distinct cell lines. Given these advantages, this protocol represents a powerful new model system for studying molecular and cellular mechanisms of yolk sac hematopoiesis and TRM differentiation. To facilitate adoption of this protocol by other laboratories, we have included a detailed, step-by-step protocol (see **Supplementary Experimental Procedures**).

*In vitro* directed differentiation models enable a variety of experimental designs that are not currently feasible *in vivo*, including large-scale genetic or chemical screens, live imaging, and biochemical studies that require large numbers of cells. Our improved *in vitro* model of mouse yolk sac hematopoiesis will thus allow for the application of these powerful experimental approaches to dissect mechanisms of cell fate specification that have proven difficult to study *in vivo*. Furthermore, using mouse EpiSCs means it is possible to take advantage of the extensive mouse genetic tools that have been developed to study yolk sac hematopoiesis, including knockout mouse lines and reporter alleles. Unlike human PSC models, findings from mouse PSC models can also be rigorously evaluated *in vivo* under controlled conditions. Given these advantages, we believe that this new mouse EpiSC model for studying yolk sac hematopoiesis and TRM differentiation can complement and empower existing model systems and will thus be of broad interest.

There is significant interest in incorporating TRMs into various tissue organoid models to better recapitulate the cellular complexity of tissues *in vivo* and to examine the crosstalk between TRMs and their host tissues ^33^. We show that mouse TRMs generated from mouse EpiSCs can integrate into developing mouse cerebral cortical organoids, generating the first chimeric neuro-immune organoid model derived completely from mouse PSCs. In the future, it will be interesting to use this system to model mechanisms of microglial development, neuroinflammation, and neurodegenerative disease. With additional development, this tractable *in vitro* model could also serve as a scalable platform for genetic or chemical screening ^80–82^.

Perhaps the most translationally relevant application of our system is the adoptive transfer of *in vitro*-derived cells into *in vivo* tissue niches. This approach serves as a powerful platform to investigate disease pathogenesis and test therapeutic efficacy *in vivo*. For instance, transferring TRMs into mouse brains or lungs could assess their ability to rescue phenotypes in models of Alzheimer’s disease or respiratory disease. Unlike human cell transplantation into mice, which requires complex immunosuppression techniques to prevent xenogeneic rejection ^83,84^, mouse-to-mouse adoptive transfers bypass this requirement when cells are matched to the host strain. This technical simplicity allows for more straightforward experimental designs and reproducible outcomes, offering new opportunities for studying therapeutic interventions.

### Limitations of the Current Study

While this new protocol can generate PHM and EMPs with reasonable efficiency, off-target mesodermal populations are still produced. Further optimization will be necessary to improve PHM and EMP yields. In addition, we have not characterized the differentiation of EMPs into other lineages (megakaryocytes, granulocytes, mast cells) ^67^.

Although we observed that culturing immature TRMs in B27-TIC media increased expression of some microglia-enriched genes, this *in vitro* model does not fully recapitulate the gene expression signatures of microglia *in vivo* (consistent with previous studies) ^72^. Achieving fully mature TRM identities *in vitro* will require exposure to additional niche-specific cues, including organ-specific extracellular matrix components or biomechanical cues ^85^.

Finally, while our adoptive transfer studies confirm that *in vitro*-derived TRMs can integrate into host tissues, the long-term stability and functional equivalence of these cells compared to endogenous TRMs will require further investigation. Future longitudinal studies tracking engrafted TRMs in vivo will be essential for assessing their durability and evaluating their therapeutic potential.

## Supporting information

Key Resources Table and Supplementary Tables

Supplementary Experimental Procedures

## ACKNOWLEDGEMENTS

We would like to thank our colleagues in the Vierbuchen and Perry laboratories for helpful discussions and advice throughout the course of the study. We would also like to thank Tomi Lazarov and Laura Santini for providing helpful comments on the manuscript.

This study would not have been possible without the core facilities at MSKCC. We would like to specifically thank members of the Flow Cytometry Core facility and the Integrated Genomics Operation Core for their advice and assistance with experiments and data analysis throughout the course of this project.

Some figure panels were created using the commercial version of Biorender.

## AUTHOR CONTRIBUTIONS

*Conceptualization*: A.K.B., J.S.A.P., T.V.

*Methodology*: A.K.B., R.V., Z.W., J.E.R.P., K.F., M.I., S.D, S.C.D., Y.L., A.T.

*Formal Analysis*: A.K.B., R.V., D.M.C., Y.L.

*Investigation*: A.K.B., R.V.

*Resources*: J.S.A.P., T.V.

*Data Curation*: A.K.B., R.V.

*Writing – original draft*: A.K.B., J.S.A.P., T.V.

*Writing – review and editing*: A.K.B., J.S.A.P., T.V.

*Supervision*: J.S.A.P., T.V.

*Project Administration*: J.S.A.P., T.V.

*Funding Acquisition*: J.S.A.P., T.V.

## DATA AVAILABILITY

Source data and processed files are available in the Gene Expression Omnibus under the accession number GSE296070.

## MATERIALS AND METHODS

### Differentiation of EpiSCs into immature TRMs

Mouse EpiSCs were generated from ESCs as described in previous work from our lab ^50^. The three EpiSC cell lines used were 1) C57BL/6J X 129S1/SvImJ F1 hybrid genetic background (B6/129), 2) DO-202 ^86^, and 3) DO-223 ^86^. EpiSCs were karyotyped to assess chromosomal integrity and tested for mycoplasma to rule out bacterial contamination. EpiSCs were cultured on gelatin-coated dishes with irradiated mouse embryonic fibroblasts in N2B27 media (50% DMEM-F12, 50% Neurobasal medium supplemented with 0.5% N2 supplement, 1% B27 supplement without vitamin A, 2 mM GlutaMAX, 1% penicillin-streptomycin, 0.1% 2-mercaptoethanol) supplemented with 20 ng/mL Activin A, 12.5 ng/mL heat-stable bFGF (FGF2), and 175 nM NVP-TNKS656. Media was changed daily, and cells were passaged every ∼48 hours at a 1:6 ratio after dissociation with Accutase into small clumps of 3-5 cells. All cells were maintained at 37°C in 5% CO_2_ under normoxic conditions.

EpiSCs between passages 8-12 were cultured to confluency and detached from the fibroblast layer using 0.1 U/μL collagenase IV, followed by resuspension in Accutase for 1-2 minutes to obtain a single-cell suspension. Plates for differentiation were sequentially coated first with 16.7 µg/mL Fibronectin in PBS^-/-^, washed twice with PBS^-/-^, and then coated with 10 μg/mL Laminin521 in PBS^+/+^ overnight. Cells were plated at a density of 60,000 cells/cm^2^ in Lo FABC-R media, which consisted of chemically defined medium 2 (CDM2) (See supplementary Table 1 for recipe) supplemented with 5 ng/mL FGF2, 30 ng/mL BMP4, 5 ng/mL Activin A, 3 µM CHIR 99021, and 50 nM Chroman-1 and 5 µM Emricasan (See Supplementary Table 1 for detailed information on how to prepare media) .

After 24 hours, the medium was switched to Hi FAB media (CDM supplemented with 10 ng/mL FGF2, 10 ng/mL Activin A, and 40 ng/mL BMP4). On day 2, media was replaced with VF media (Essential 6 medium containing 15 ng/mL VEGF and 5 ng/mL FGF2). VF media was refreshed at the end of day 3. On day 4, cells were cultured in VFSI media (Essential 6 medium supplemented with 15 ng/mL VEGF, 5 ng/mL FGF2, 200 ng/mL SCF, and 20 ng/mL IL-6). On day 5, cells were transferred to SITI media (Essential 6 medium containing 100 ng/mL SCF, 10 ng/mL IL-6, 30 ng/mL TPO, and 30 ng/mL IL-3), with daily media changes until day 7 or 8. Cells were never washed between media changes.

On day 7 or 8, cells were dissociated into single cells with Accutase and replated in macrophage media at a density of 60,000–120,000 cells per cm^2^ on Matrigel-coated plates. Macrophage media consisted of 75% IMDM with GlutaMAX, 25% F12 with GlutaMAX, 1:100 B27 supplement without vitamin A, 100 μg/mL penicillin-streptomycin, 100 ng/mL mouse M-CSF, and 100 ng/mL mouse IL-34. Macrophage media was changed three times a week, and cells were passaged every 7-8 days by simply pipetting to detach them from the plates and replating at 60,000 cells per cm^2^.

Additional details about reagents and differentiation media compositions are provided in Supplementary Table 1 and the Key Resources Table.

### Fluorescence-Activated Cell Sorting (FACS) analysis

For 2D cultured cells, cell dissociation was performed with Accutase for 1-2 minutes, followed by washing in FACS buffer (1% BSA, 5 mM EDTA). Organoid cells were dissociated using Accutase containing 30 μg/mL DNase I for 30-45 minutes. After dissociation, cells were washed and resuspended in FACS buffer with the appropriate antibodies.

Day 16+ macrophages were incubated with Fc block (unconjugated mouse anti-CD16/32 Bio X Cell, 1:50) for 10 minutes to prevent nonspecific Fc receptor binding before antibody staining. Cells were then washed, resuspended in FACS buffer, and filtered through 40 μm filters prior to analysis on an Attune Nxt flow cytometer. Data was analyzed using FlowJo v10.10.0.

Sorting of CX_3_CR1^+^ macrophages was done at the MSKCC Flow Cytometry Core Facility using either BD FACSAria or BD FACSymphony S6 cell sorters.

FACS antibodies and dilutions used were as follows:

Day 2 cells: KDR-PE (BioLegend, 1:200), PDGFRα-PE/Cy7 (BioLegend, 1:200), CXCR4-Alexa Fluor 647 (BioLegend, 1:200)

Day 8 cells: Kit-BV421 (BD Biosciences, 1:200), CD41-FITC (BioLegend, 1:200), CD16/32-BV711 (BioLegend, 1:200), CD45-Alexa Fluor 700 (BioLegend, 1:200)

Day 16+ macrophages: unconjugated anti-mouse CD16/32, CX_3_CR1-BV785 (BioLegend, 1:200), CD11b-FITC (BioLegend, 1:200), CD45-Alexa Fluor 700 (BioLegend, 1:200), P2RY12-PE (BioLegend, 1:200)

Detailed information on all FACS antibodies is available in Supplementary Table 2.

### Immunostaining of 2D cells

Cells were washed twice with PBS-/- and fixed with 4% paraformaldehyde (PFA) for 15 minutes at room temperature. After fixation, cells were rinsed twice with PBS-/- and permeabilized with PBS-/- containing 1% Triton-X for 10 minutes at room temperature. Cells were blocked for 30 minutes with PBS-/- with 0.3% Triton-X and 5% BSA. Primary antibodies, diluted in PBS containing 0.1% Triton-X (PBST) with 1% BSA, were then added overnight at 4°C. The following primary antibodies were used: rabbit anti-Brachyury/T (Abcam, 1:1000), goat anti-Brachyury/T (R&D Systems, 1:200), goat anti-FOXF1 (R&D Systems, 1:400), rabbit anti-GATA6 (Cell Signaling Technology, 1:500), rabbit anti-VE-Cadherin (ThermoScientific, 1:200), rabbit anti-Runx1 (ThermoScientific, 1:400), goat anti-Iba1 (Novus Biologicals, 1:500), rabbit anti-Pu.1 (ThermoScientific, 1:200), rat anti-Ki-67 (Invitrogen, 1:500). Detailed information about antibodies can be found in Supplementary Table 3.

After overnight incubation with primary antibodies, cells were washed three times with PBST for 10 minutes each. Secondary antibodies, diluted 1:500 in PBST with 1% FBS, were added for 2 hours at room temperature. After three additional 10-minute washes with PBST, cells were stained with DAPI (1 μM in PBST) for 15 minutes at room temperature. Cells were then rinsed with PBST and imaged using a Leica Dmi8 microscope.

### Differentiation of immature TRMs into microglia-like cells

After 3-4 days of proliferation in macrophage media, around day 11, cells were transitioned to microglia media without additional passaging. Two formulations of microglia media were tested: the first, TIC media, is an established serum-free medium for culturing primary mouse and rat microglia^72^. This media contains DMEM-F12, 2 mM GlutaMAX, 100 μg/mL penicillin-streptomycin, 1X Insulin-Transferrin-Selenium, and 1 mg/mL N-acetyl cysteine (NAc), supplemented with 2 μg/mL TGF-β, 100 ng/mL IL-34, 1.5 μg/mL cholesterol, 0.1 mg/mL oleic acid, 0.001 μg/mL gondoic acid, and 10 μg/mL heparan sulfate.

The second formulation, a modified TIC media we refer to as B27-TIC, comprised of a base of 75% IMDM with GlutaMAX, 25% F12 with GlutaMAX, and 1:100 B27 supplement without vitamin A, supplemented identically to TIC media. After 5-6 days in TIC media, cells were split and replated at a density of 120,000 cells/cm^2^ on Matrigel- or collagen IV-coated plates, with collagen IV-coated plates yielding more highly ramified cells. Cells in B27-TIC media adhered better to coated tissue culture plates.

Additional details about reagents and differentiation media compositions are provided in the KEY RESOURCES TABLE and Supplementary Table 1.

### Primary microglia isolation and culture

Neonatal brains were collected from P4 WT C57BL/6J mice and dissociated using the Miltenyi gentleMACS Octo Dissociator with Heaters, brain dissociation kits (Miltenyi Biotec, 130-096-427, 130-107-677, 130-092-628), and gentleMACS C Tubes (Miltenyi Biotec, 130-093-237), following the manufacturer’s instructions. Microglia were then purified using CD11b MicroBeads (Miltenyi Biotec, 130-097-142), LS Columns (Miltenyi Biotec, 130-042-401), and the QuadroMACS separator (Miltenyi Biotec, 130-091-051) following manufacturer’s instructions.

Purified microglia were cultured in B27-TIC media. Cells were maintained under standard conditions at 37°C in 5% CO₂ within a normoxic tissue culture incubator.

### Bulk RNA sequencing

Approximately 500,000 -1,000,000 CX3CR1^+^ immature macrophages or microglia-like cells were sorted from three distinct mouse cell lines. Sorted cells were pelleted by centrifugation, frozen, and sent to the Memorial Sloan Kettering Cancer Center (MSKCC) Integrated Genomics Operation Core (IGO) for RNA extraction and quality control. The core performed RNA-polyA sequencing, generating 30–40 million reads per sample.

Resulting FASTQ files were processed using Salmon^87^ for transcript mapping and quantification against the mm10 mouse genome. Downstream analyses were conducted in RStudio, using the packages DESeq2, pcaExplorer, and pheatmap^88,89^. Statistical significance was assessed using the Wald test and corrected with the Benjamini–Hochberg method.

### Phagocytosis assays

Macrophages were first plated on collagen-IV-coated 8-well glass chamber slides (Nunc, 155409 for live imaging, 154534 for fixed imaging) and allowed to adhere overnight before addition of phagocytosis targets.

For the synaptosome phagocytosis assay, synaptosomes were isolated from wild-type adult mice using the Syn-PER Synaptic Protein Extraction Reagent (ThermoScientific, 87793) following the manufacturer’s protocol. Synaptosomes were stained with CypHer 5E (GE Life Sciences, PA15401) via agitation, washing, and sonication. A BCA assay confirmed the staining concentration (10 mM CypHer 5E/50 µL). Stained synaptosomes (20 µL) were added to macrophages, and live imaging was performed for 45 minutes.

For the apoptotic neuron phagocytosis assay, apoptosis was induced in Neuro-2a cells (ATCC, CCL-131) using 0.25 µM staurosporine (Cayman Chemical, 81590) for 12-15 hours. Apoptotic neurons were stained with 1 μM CypHer 5E in serum-free HBSS for 45 minutes at 37°C, then washed in DMEM for 20 minutes at 37°C. These neurons were resuspended in either immature TRM media or B27-TIC and incubated with macrophages at a 1:1 ratio for 45 minutes during live imaging.

For apoptotic Jurkat cells, apoptosis was induced with 150 mJ/cm^2^ UVC irradiation followed by a 4-hour incubation at 37°C and 5% CO₂. Cells were stained with 1 μM CypHer 5E in serum-free HBSS for 45 minutes at 37°C, washed, and incubated in serum-complete medium for 25 minutes. Macrophages were fed apoptotic Jurkat cells at a 1:1 ratio for 45 minutes. Post-incubation, cells were washed, fixed with 4% PFA, and permeabilized with 0.3% Triton-X for 20 minutes. After blocking with 4% goat serum and 1% BSA, cells were incubated overnight at 4°C with primary antibodies: goat anti-Iba1 (FUJIFILM Wako Chemicals, 1:500) and rabbit anti-CD68 (ThermoScientific, 1:250). Secondary antibodies (1:500) were applied for 1 hour at room temperature. Cells were washed, mounted with Prolong Gold, and imaged the next day.

For zymosan phagocytosis, AlexaFluor 594-conjugated zymosan particles (ThermoScientific, Z23374) were added to macrophages at a 1:1 ratio, followed by the same fixation and staining protocol used for Jurkat cell phagocytosis.

Live imaging of phagocytosis was conducted using time-lapse confocal microscopy on a Zeiss Axio Observer.Z1-7 with a Z PIEZO stage, heating/gas-controlled insert, Zeiss 20x PlanApo objective, LSM 980 with Airyscan.2 multiplex detector. Images were captured every 30 seconds. Fixed cell imaging was also performed on the same system. All data was analyzed with Zen Blue software (Zeiss) and FIJI.

### Generation of cerebral cortical organoids and TRM transplantation

Cortical organoids devoid of macrophages were first generated from EpiSCs using a previously described protocol^55^. Briefly, EpiSCs were seeded at 1,000 cells per microwell in an AggreWell 400 plate (34415, STEMCELL Technologies) using EB formation media consisting of 50 nM chroman-1, 5 μM emricasan, 400 nM LDN-193189, 10 μM SB-431542, 1 μM cyclopamine, and 100 nM LGK-974 in N2B27 (B27 without vitamin A) media. After 24 hours, EBs were transferred to the same media, but without chroman-1 and emricasan, and embedded in Matrigel. To prepare the Matrigel mix, 66.7 μL of EBs in media were combined with 100 μL of Matrigel using wide-bore tips, then plated in a 6-well plate without touching well edges and incubated at 37°C for 30 minutes. Following incubation, 3 mL of warm media was added on top. On day 2, media was replaced with plain N2B27 containing 1 μM cyclopamine and 1% KSR for 24 hours. On day 3, media was replaced with N2B27 supplemented with 1 μM cyclopamine, 3 μM CHIR99201, and 1% KSR. On day 4, organoids were recovered from Matrigel using Cell Recovery Solution (Corning, 354270). Wells were washed with PBS, and 2–3 mL of Cell Recovery Solution was added, ensuring the Matrigel dome was lifted for efficient organoid retrieval. Plates were incubated at 4°C for 30 minutes, followed by two washes in organoid media (N2B27 with 20% KSR). Finally, cortical organoids were transferred to Petri dishes and maintained in N2B27 with 20% KSR while shaking at 65 rpm (Infors HT, Celltron benchtop shaker). Cortical organoid media (N2B27 with 20% KSR) was changed every other day until day 10, when immature TRMs were added.

To integrate immature TRMs, a single organoid was transferred to an ultra-low attachment U-bottom 96-well plate. Day 20-28 immature TRMs (50,000 cells per organoid) were added in cortical organoid media supplemented with 100 ng/mL IL-34 and 2 ng/mL TGF-β. The immature TRMs were allowed to spontaneously migrate into the organoids overnight at 37°C, 5% CO₂.

The organoids were then pooled and transferred to 10 cm plates, which were placed on the orbital shaker. Any remaining immature TRMs from the 96-well plate were added to the 10 cm plate to maximize integration. These neuroimmune organoids were cultured for an additional 2-3 weeks in organoid media supplemented with IL-34 (100 ng/mL) and TGF-β (2 ng/mL), with media changes twice a week. Organoids without immature TRMs were cultured in parallel as controls.

### Immunostaining of organoids

Organoids were collected in 1% BSA-coated tubes, washed twice with PBS-/- and fixed in 4% paraformaldehyde for 45 minutes at 4°C. After fixation, samples were washed twice with PBS-/- and permeabilized using 0.5% Triton X-100 for 15 minutes at 4°C. Blocking was carried out for 15 minutes in Organoid Washing Solution (OWS; adapted from Dekkers et al., 2019), composed of 0.2% Triton X-100, 0.02% SDS, and 0.2% BSA in PBS-/-, at 4°C.

Organoids were incubated overnight at 4°C with primary antibodies diluted in OWS. The following day, they were washed three times with OWS for 2 hours each under constant shaking, followed by overnight incubation with secondary antibodies and DAPI (1 µM) in OWB at 4°C. The organoids were then washed three times with OWB over a total of 6 hours while shaking. Finally, organoids were mounted in DeepClear solution (CelExplorer Labs, DC-201) and imaged using a Leica SP8 confocal microscope. Image analysis was performed with Imaris software.

### Adoptive transfer of immature TRMs into mouse lung

To deplete endogenous alveolar macrophages, 8-week-old CD45.1 congenic male and female mice were anesthetized with isoflurane and administered 50 μL of clodronate liposomes (Fisher Scientific, NC0337390) intratracheally. After 3 days, mice were anesthetized again and given 300,000 immature TRMs in 50 μL of sterile PBS intratracheally.

On days 3 and 7 post-transfer, mice were euthanized and perfused with a 21G needle containing 40 mL of cold PBS-/-(Corning, 21-040-CM) and 5 mM EDTA (Invitrogen, 15575-038). One lung per mouse was dissected, minced into small fragments, and digested for 30 minutes at 37°C in enzyme solution comprising PBS with 160 Wünsch U/mL collagenase D (Sigma, 11088866001), 5% heat-inactivated FBS (Sigma, 12306C), and 10 mM HEPES (Gibco, 15630-080) (referred to as PFH buffer). The reaction was stopped with 10 mM EDTA for 5 minutes.

Digested lung tissue was further dissociated mechanically using a 70 μm cell strainer and an 18G needle with 3-mL syringe plunger in 6-well plates containing 4 mL of cold PFH buffer. The resulting single-cell suspensions were transferred to 15-mL tubes and pelleted by centrifugation at 1500 rpm for 5 minutes at 4°C. Pellets were resuspended in 38% isotonic Percoll (10% 10X PBS, 90% Percoll; GE Healthcare, 17-0891-01) in PFH buffer to gradient separate leukocytes by centrifugation at 2000 rpm for 30 minutes at room temperature with no brake.

To evaluate the differentiation of transferred immature TRMs into mature alveolar macrophages, lung cell suspensions were incubated with Fc block for 10 minutes, followed by staining with the following antibodies: CD45.2-AlexaFluor 700 (BioLegend, 1:400), CD11b-PE (Thermo Scientific, 1:800), CD11c-APC (BioLegend, 1:800), and SiglecF-BV421 (BioLegend, 1:400).

Detailed information on antibodies is provided in Supplementary Table 2.

### Statistical analysis and reproducibility

Statistical tests were performed using GraphPad Prism 10 or Rstudio. Experiments with statistical analyses were performed with a minimum of three independent differentiation experiments (technical replicates) or biological replicates (different cell lines). For experiments involving live mice, each data point represents results from a single mouse (lung adoptive transfer) or from cells pooled from 4 different mice (*ex vivo* microglia cultures).

## SUPPLEMENTARY INFORMATION

### Key Resources Table

**Supplementary Experimental Procedures:** Step-by-step protocol for the directed differentiation of mouse EpiSCs into immature TRMs and microglia-like cells **Supplementary Table 1:** 2D Differentiation media compositions

**Supplementary Table 2**: List of FACS antibodies

**Supplementary Table 3**: List of IF antibodies

**Supplementary Video**: Phagocytosis of CypHer 5E-labelled synaptosomes by immature TRMs

---

**Supplementary Figure 1:**
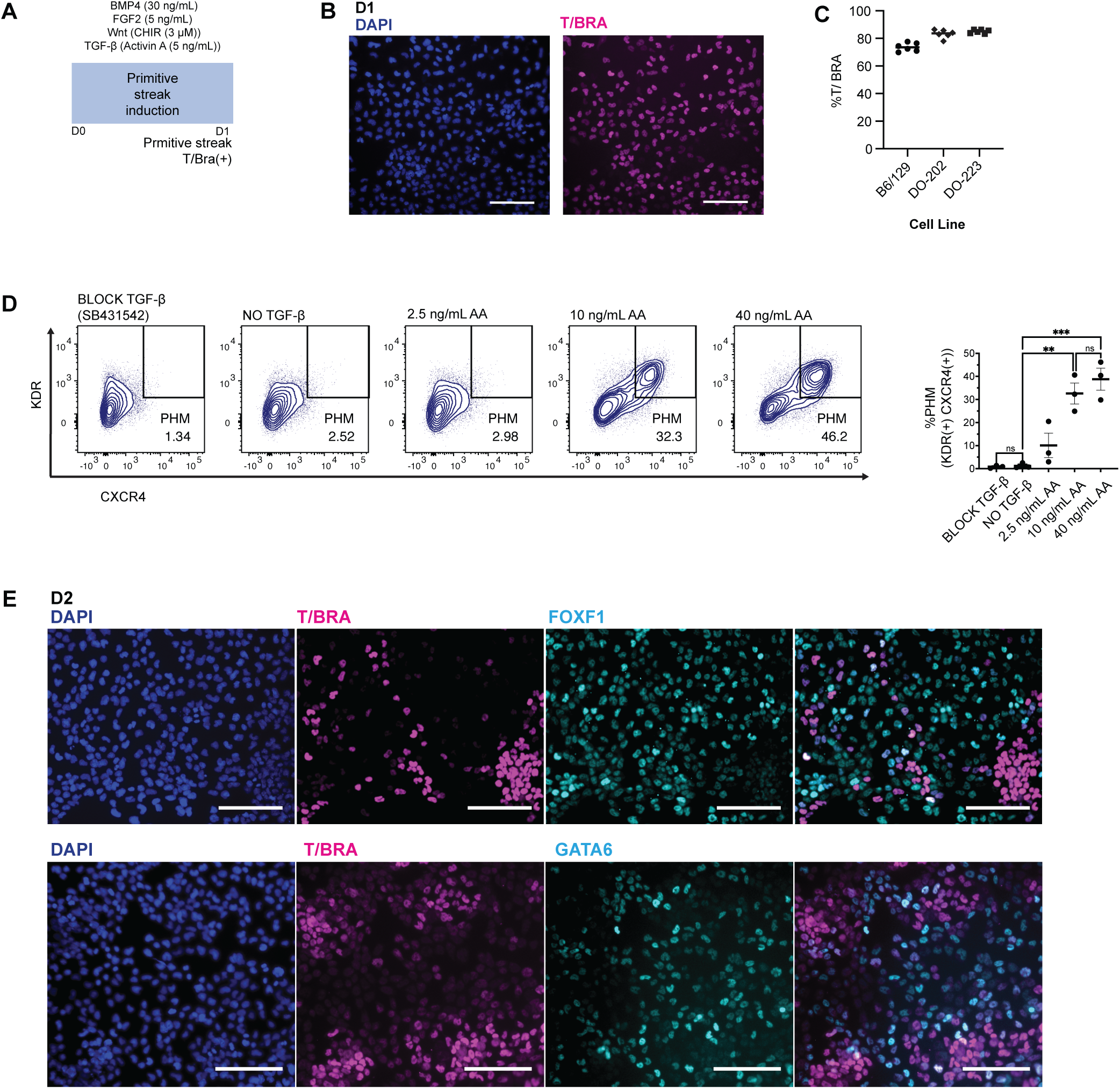
Identification of conditions for generating yolk sac pre-hematopoietic mesoderm (PHM) from mouse EpiSCs. **A**. Summary of the first stage of the TRM differentiation protocol. **B**. Immunofluorescence staining for brachyury (T) was used to evaluate the percentage of cells that have acquired a primitive streak-like identity at the end of stage 1 (d1). **C**. Quantification of Brachyury (T) positive cells at the end of stage 1 across 3 distinct EpiSC lines (B6/129, DO-202, DO-223). Three independent differentiation experiments were performed for each EpiSC line, with two technical replicates per cell line. **D**. Representative flow cytometry plots showing D2 KDR(+) CXCR4 (+) yield across TGF-β conditions (left). Summary of flow cytometry data across 3 independent differentiation experiments (right). Significance was determined using ordinary one-way ANOVA, corrected for multiple comparisons with Tukey test (not significant [ns] P > 0.05; **P < 0.01; ***P < 0.001). **E**. Immunofluorescence staining for additional markers of PHM (FOXF1, GATA6) in differentiating cultures at the end of Stage 2 (D2). Scale bars = 100 µm.

**Supplementary Figure 2:**
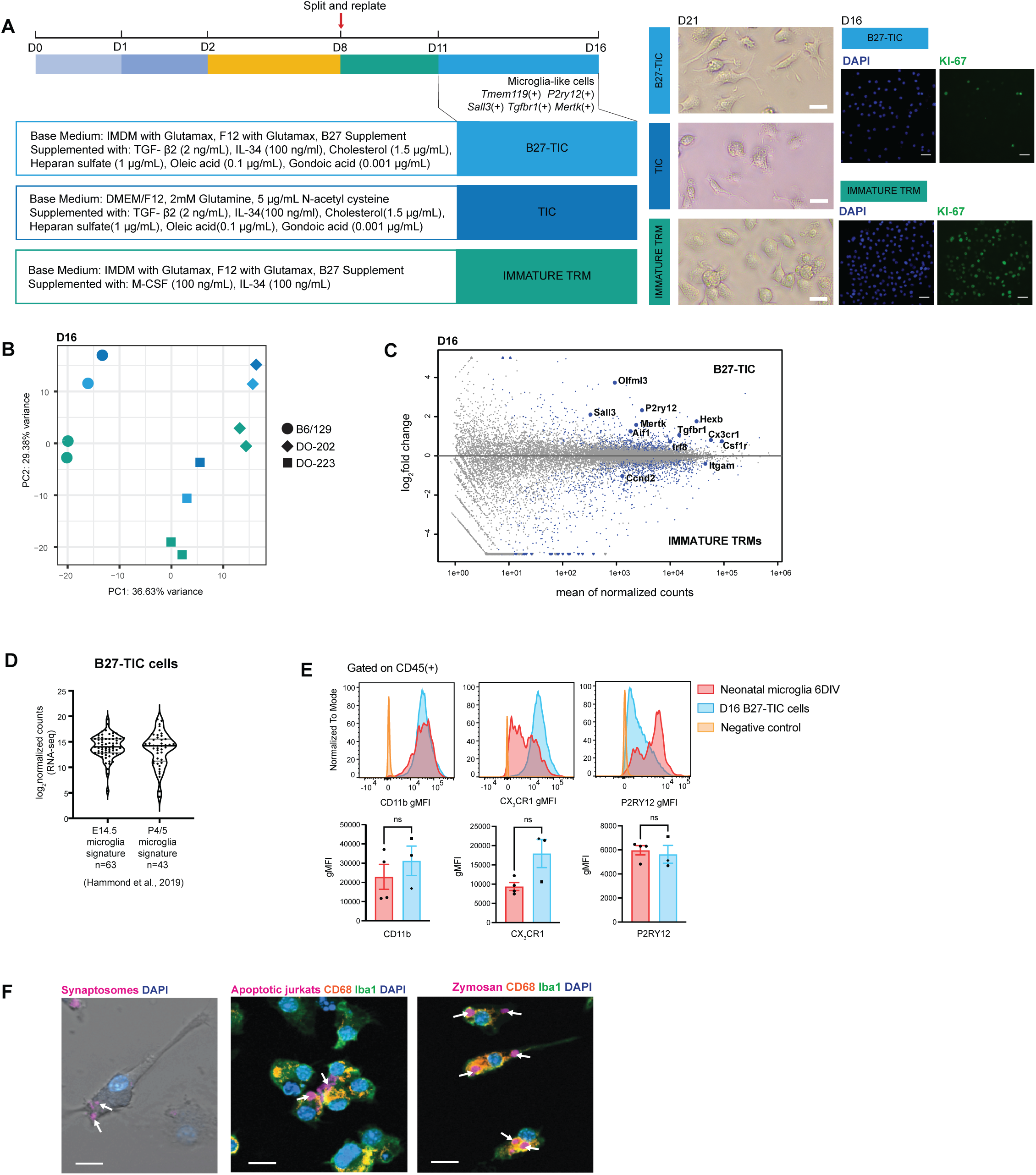
Characterization of TRMs cultured in *ex vivo* microglia maintenance conditions. **A**. Overview of different media conditions for culturing immature TRMs (left). Bright field images of TRMs in each condition (right) and immunofluorescence staining for cell proliferation (Ki-67) (far right). **B**. Principal component analysis (PCA) plot of RNA-seq data from D16 TRMs cultured in each of the three media conditions (color coded based on [A]). **C**. MA plot comparing gene expression of D16 immature TRMs and TRMs cultured in B27-TIC. A panel of genes relevant for immature TRMs and microglia are annotated. Statistical significance was calculated using a Wald test and corrected using the Benjamini-Hochberg method; gray dots P > 0.1; blue dots P < 0.1 **D**. Violin plots of gene expression (RNA-seq) in D16 B27-TIC cells. Gene signatures are derived from microglia isolated at E14.5 or P4/P5 (Hammond et al., 2019). Each dot represents the average expression level of gene (n = 3 biological replicates). **E**. Representative geometric mean fluorescence intensity (gMFI) plots comparing microglia surface marker profiles of primary microglia and B27-TIC microglia-like cells; gated for CD45(+) expression (top row). Quantification of gMFI across biological replicates (bottom row). Statistical significance was determined using Welch’s t-test (ns P > 0.05). **F**. Confocal imaging of B27-TIC TRMs (D20) phagocytosing pH-sensitive CypHer 5E-labeled synaptosomes (left). Immunofluorescence staining for pan-macrophage marker CD68 and IBA1 in fixed B27-TIC TRMs that were exposed to CypHer 5E-labeled apoptotic Jurkat T cells or AlexaFluor 594-conjugated zymosan beads. Arrows indicate engulfed targets (right). Scale bars = 25 µm. DIV = days *in vitro*.

**Supplementary Figure 3:**
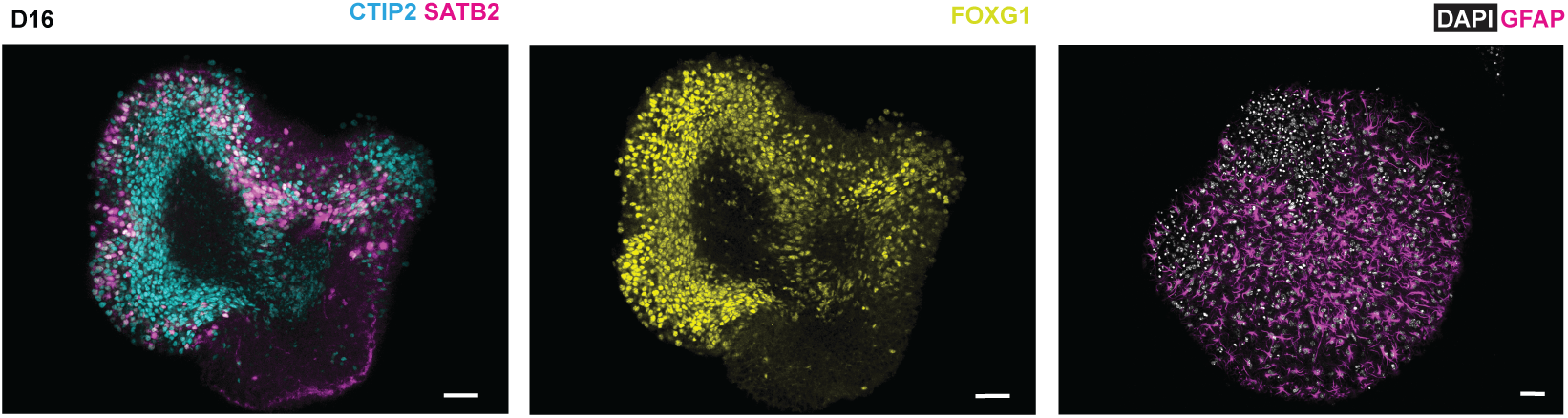
Immature TRMs integrate into developing cortical organoids and adult mouse lung niche. Immunofluorescence staining of D16 cortical organoids for markers of deep-layer cortical neurons (CTIP2), upper-layer neurons (SATB2), telencephalic progenitors (FOXG1) and astrocytes (GFAP). Scale bars = 30 µm

