## Supplementary material for "Modeling tissue-resident macrophage development from mouse pluripotent stem cells": Key Resources Table and Supplementary Tables

| Reagent/ Resource | Company | Catalog Number |
|---|---|---|
| Accutase | Gibco | Cat#A1110501 |
| Activin A | Peprotech | Cat#120-14P |
| Adult Brain Dissociation Kit, mouse and rat | Miltenyi Biotec | Cat#130-107-677 |
| AggreWell™400 | STEMCELL Technologies | Cat#34415 |
| B27 supplement (without vitamin A) | Gibco | Cat#12587010 |
| Bovine Serum Albumin (BSA) | Sigma-Aldrich | Cat#A2153 |
| CD11b MicroBeads | Miltenyi Biotec | Cat#130-097-142 |
| Cell recovery solution | Corning | Cat#354270 |
| Chemically Defined Lipid Concentrate | Gibco | Cat#11905031 |
| CHIR99201 | Tocris Bioscience | Cat#252917-06-9 |
| Cholesterol (ovine) | Avanti Polar Lipids | Cat#700000P |
| Chroman 1 | MedChem Express | Cat#HY-15392 |
| Clodronate Liposomes | Fisher Scientific | Cat#NC0337390 |
| Collagen IV, Mouse | Corning | Cat#354233 |
| Collagenase D | Sigma-Aldrich | Cat#11088866001 |
| Collagenase Type IV | Gibco | Cat#17104019 |
| Cyclopamine | Cayman Chemical Company | Cat#11321 |
| CypHer 5E | GE Life Sciences | Cat#PA15401 |
| DeepClear | Celexplorer | Cat#DC-201 |
| DMEM | Thermo Fisher Scientific | Cat#11965118 |
| DMEM/F12 medium | Gibco | Cat#11320082 |
| DNase I | Zymo Research | Cat#E1010 |
| EDTA (0.5 M), pH 8.0, RNase-free | Invitrogen | Cat#AM9261 |
| Emricasan | Selleck Chemicals | Cat#S7775 |
| gentleMACS C Tubes | Miltenyi Biotec | Cat#130-093-237 |
| gentleMACS™ Octo Dissociator with Heaters | Miltenyi Biotec | Cat#130-096-427 |
| Glutamax | Gibco | Cat#35050061 |
| Gondoic Acid (11(Z)-Eicosenoic Acid) | Cayman Chemical Company | Cat#20606 |
| Ham's F12 Nutrient Mix, GlutaMAX Supplement | Thermo Fisher Scientific | Cat#31-765-035 |
| Hbss | Thermo Scientific | Cat#14170112 |
| Heat stable recombinant human bFGF (FGF2) | Gibco | Cat#PHG0360 |
| Heat-Inactivated FBS | Sigma-Aldrich | Cat#12306C |
| Heparan Sulfate | Amsbio | Cat#AMS.GAG-HS01 |
| HEPES | Gibco | Cat#15630080 |
| Human Fibronectin | Sigma-Aldrich | Cat#FC010 |
| IMDM, GlutaMAX Supplement, HEPES | Thermo Fisher Scientific | Cat#31-980-030 |
| Insulin | Sigma-Aldrich | Cat#91077C |
| Insulin-Transferrin-Selenium | Thermo Scientific | Cat#41400045 |
| KnockOut™ Serum Replacement (KSR) | Thermo Fisher Scientific | Cat#10828028 |
| LDN-193189 | Reprocell | Cat#04-0074 |

| | | |
|---|---|---|
| LGK-974 | Selleck Chemicals | Cat#S7143 |
| LS Columns | Miltenyi Biotec | Cat#130-042-401 |
| Matrigel | Corning | Cat#354230 |
| Monothioglycerol | Sigma-Aldrich | Cat#M6145 |
| N-Acetyl-L-cysteine | Sigma-Aldrich | Cat#A9165-25G |
| N2 supplement | Gibco | Cat#17502048 |
| Neural Tissue Dissociation Kit (P) | Miltenyi Biotec | Cat#130-092-628 |
| Neuro-2a cells | ATCC | Cat#CCL-131 |
| Neurobasal medium | Gibco | Cat#21103049 |
| Normal Goat Serum | Jackson ImmunoResearch | Cat#005-000-121 |
| NVP-TNKS656 | Selleck Chemicals | Cat#S7238 |
| Oleic Acid | Cayman Chemical Company | Cat#90260 |
| Optiferrin (recombinant human transferrin) | Invitria | Cat#777TRF029 |
| Penicillin/streptomycin (100X) | Gibco | Cat#15140122 |
| Percoll | GE Healthcare | Cat#17-0891-01 |
| Polyvinyl alcohol | Sigma-Aldrich | Cat#341584 |
| ProLong Gold Antifade Mountant | Fisher Scientific | Cat#P36934 |
| QuadroMACS separator | Miltenyi Biotec | Cat#130-091-051 |
| Recombinant Human BMP-4 Protein | R&D Systems | Cat#314-BP |
| Recombinant Human IL-3 | PeproTech | Cat#200-03 |
| Recombinant Human IL-6 Protein | R&D Systems | Cat#206-IL |
| Recombinant Human SCF | PeproTech | Cat#300-07 |
| Recombinant Human TGF-β2 | PeproTech | Cat#100-35B |
| Recombinant Human TPO | PeproTech | Cat#300-18 |
| Recombinant Human VEGF 165 Protein | R&D Systems | Cat#293-VE |
| Recombinant Mouse IL-34 | BioLegend | Cat#577606 |
| Recombinant Murine M-CSF | PeproTech | Cat#315-02 |
| rhLaminin-521 | Gibco | Cat#A29249 |
| SB 431542 | Tocris | Cat#1614 |
| Sodium Dodecyl Sulfate (SDS) | Fisher Scientific | Cat#BP24361 |
| Staurosporine | Cayman Chemical Company | Cat#81590 |
| Synaptic Protein Extraction Reagent (Syn-PER) | Thermo Scientific | Cat#87793 |
| Triton X-100 | Sigma-Aldrich | Cat#9002-93-1 |
| Ultra-Low Attachment Microplate | Corning | Cat#7007 |
| 16% Formaldehyde (w/v), Methanol free | Thermo Scientific | Cat#28908 |
| 2-Mercaptoethanol | Gibco | Cat#21985023 |
| 8-well Chamber Slide w/ removable wells | Thermo Scientific | Cat#154534 |
| 8-well Chambered Coverglass w/ non-removable wells | Thermo Scientific | Cat#155409 |

**Supplementary Table 1: 2D Differentiation media compositions**

| Medium | Recipe | Comments |
|---|---|---|
| CDM2 | 50% IMDM (with Glutamax), 50% F12 (with Glutamax), 1X Chemically Defined Lipid Concentrate, 450 µM Monothioglycerol, 1 mg/mL Polyvinyl Alcohol, 15 µg/mL Optiferrin, 0.7 µg/mL Insulin, 1:100 Penicillin-Streptomycin | D0-D2 base medium.<br>Can store at 4°C for up to 1 month. |
| Lo FABC-CE | CDM2 containing 5 ng/mL FGF2, 5 ng/mL Activin A, 30 ng/mL BMP4, 3 µM CHIR99201, 50 nM Chroman-1, 5 µM Emricasan | D0-D1 differentiation medium. Prepare no more than 2 days before use. |
| Hi FAB | CDM2 containing 10 ng/mL FGF2, 10 ng/mL Activin A, 40 ng/mL BMP4 | D1-D2 differentiation medium. Prepare no more than 2 days before use. |
| Essential 6 Medium | Essential 6 medium containing 1:100 Penicillin-Streptomycin | D2-D8 base medium.<br>Can store for up to 1 month. |
| VF | Essential 6 medium containing 15 ng/mL VEGF, 5 ng/mL FGF2 | D2-D4 differentiation medium. Prepare no more than 2 days before use. |
| VFSI | Essential 6 medium containing 15 ng/mL VEGF, 5 ng/mL FGF2, 200 ng/mL SCF, 20 ng/mL IL-6 | D4-D5 differentiation medium. Prepare no more than 2 days before use. |
| SITI | Essential 6 medium containing 100 ng/mL SCF, 10 ng/mL IL-6, 30 ng/mL TPO, 30 ng/mL IL-3 | D5-D8 differentiation medium. Prepare no more than 2 days before use. |
| Macrophage base medium | 75% IMDM (with Glutamax), 25% F12 (with Glutamax), 1X B27 Supplement, 1:100 Penicillin-Streptomycin | Immature TRM and B27-TIC base medium.<br>Can be stored for up to one month at 4°C. |
| Immature TRM medium | Macrophage base medium containing 100 ng/mL mouse M-CSF and 100 ng/mL mouse IL-34 | Immature TRM differentiation and expansion medium. |
| B27-TIC | Macrophage base medium containing 2 ng/mL TGF-β2, 100 ng/mL mouse IL-34, 1.5 µg/mL ovine wool cholesterol, 10 µg/mL heparan sulfate, 0.1 µg/ml oleic acid, and 0.001 µg/ml gondoic acid | D11+ medium for differentiation of immature TRMs into microglia-like cells.<br><br>Make sure to add cholesterol to media warmed to 37 °C and do not add more than 1.5 µg/mL or it will precipitate out. Do not filter cholesterol-containing media.<br><br>Do not store for more than a week. |
| TIC base medium | DMEM/F12 containing 1:100 Penicillin-Streptomycin, 2 mM glutamine, 5 µg/mL N-acetyl cysteine, 5 µg/mL insulin, 1X Insulin-Transferrin-Selenite (ITS) | D11+ base medium for differentiation into microglia-like cells.<br>Can be stored at 4°C for up to one month. |
| TIC medium | TIC base medium containing 2 ng/mL TGF-β2, 100 ng/mL mouse IL-34, 1.5 µg/mL ovine wool cholesterol, 10 µg/mL heparan sulfate, 0.1 µg/ml oleic acid, and 0.001 µg/ml gondoic acid | D11+ medium for differentiation of immature TRMs into microglia-like cells. |

| | | |
|---|---|---|
| | | Make sure to add cholesterol to media warmed to 37 °C and do not add more than 1.5 µg/mL or it will precipitate out. Do not filter cholesterol-containing media.<br><br>Do not store for more than a week. |
| **Stock Reagent** | **Preparation details** | **Comments** |
| Fibronectin (liquid) | Stock conc: 1 mg/mL<br>Reconstitution: N/A<br>Use at: 167 µL/mL | store at 4°C |
| Laminin-521 | Stock conc: 100 µg/mL<br>Reconstitution: N/A<br>Use at: 1:10 | Thaw at 4°C and do not freeze-thaw |
| Polyvinyl Alcohol | Stock conc: 100 mg/mL<br>Reconstitution: In a 250ml or larger glass bottle, add 100mL of di$H_2O$ to 10g PVA powder, then autoclave under dry settings ror 45-60minutes.<br>Use at: 1:100<br>Storage: 4°C | Can store at 4°C for 3 months. |
| Optiferrin | Stock conc: 10 mg/mL<br>Reconstitution: Add 100mL di$H_2O$ to 1g Optiferrin powder and mix until complete homogenization.<br>Storage: -80°C | Aliquot stock and store at -80°C for up too 12 months |
| Insulin | Stock conc: 10 mg/mL<br>Reconstitution: Take 250 mg, add 25 mL diH2O. This creates a cloudy solution. Add HCl until a pH=3.0 is achieved.<br>Storage: 4°C | Store at 4°C |
| FGF2 | Stock conc: 25 µg/mL<br>Reconstitution: Reconsitute to 1g/L in di$H_2O$. Then, add PBS + 0.1% BSA to dilute to 25 µg/mL.<br>Storage: -20°C | Aliquot stock and store at -80°C for up too 12 months |
| Activin A | Stock conc: 100 µg/mL (or less)<br>Reconstitution: Reconstitute lyophilized protein to 100ug/ml in sterile $H_2O$ with 0.1% BSA.<br>Storage: -80°C | Aliquot stock and store at -80°C for up to 12 months |
| BMP4 | Stock conc: 50 µg/mL<br>Reconstitution: Reconstitute to 50 µg/mL in 4mM HCl.<br>Storage: -80°C | Aliquot stock and store at -80°C for up to 6 months |
| CHIR99201 | Stock conc: 5 mM<br>Reconstitution: Resuspend in DMSO to make 5 mM stock<br>Storage: -20°C | Aliquot stock and store at -20°C for up to 12 months |
| Chroman-1 | Stock conc: 1 mM<br>Reconstitution: Resuspend in DMSO to make 1 mM stock<br>Storage: -80°C | Aliquot stock and store at -80°C for up to 6 months |
| Emricasan | Stock conc: 50 mM | Aliquot stock and store at |

| | | |
|---|---|---|
| | Reconstitution: Resuspend in DMSO to make 50 mM stock<br>Storage: -80°C | -80°C for up to 2 years |
| VEGF | Stock conc: 100 µg/mL<br>Reconstitution: Resuspend in PBS + 0.1% BSA to make 100 µg/mL stock<br>Storage: -80°C | Aliquot stock and store at -80°C for up to 6 months |
| SCF | Stock conc: 200 µg/mL<br>Reconstitution: Resuspend in PBS + 0.1% BSA to make 200 µg/mL stock<br>Storage: -80°C | Aliquot stock and store at -80°C for up to 6 months |
| IL-6 | Stock conc: 100 µg/mL<br>Reconstitution: Resuspend in PBS + 0.1% BSA to make 100 µg/mL stock<br>Storage: -80°C | Aliquot stock and store at -80°C for up to 6 months |
| TPO | Stock conc: 100 µg/mL<br>Reconstitution: Resuspend in PBS + 0.1% BSA to make 100 µg/mL stock<br>Storage: -80°C | Aliquot stock and store at -80°C for up to 6 months |
| IL-3 | Stock conc: 100 µg/mL<br>Reconstitution: Resuspend in PBS + 0.1% BSA to make 100 µg/mL stock<br>Storage: -80°C | Aliquot stock and store at -80°C for up to 6 months |
| M-CSF | Stock conc: 100 µg/mL<br>Reconstitution: Resuspend in sterile PBS to make 100 µg/mL stock<br>Storage: -80°C | Aliquot stock and store at -80°C for up to 6 months |
| IL-34 | Stock conc: 100 µg/mL<br>Reconstitution: comes already reconstituted at 100 µg/mL. If 100 µg is bought, be sure to bring up total volume to 1mL.<br>Storage: -80°C | Aliquot stock and store at -80°C for up to 6 months |
| TGF-β2 | Stock conc: 2 µg/mL<br>Reconstitution: Resuspend in sterile PBS to make 2 µg/mL stock<br>Storage: -20°C | Aliquot stock and store at -20°C for up to 6 months |
| Cholesterol (ovine) | Stock conc: 1.5 mg/mL<br>Reconstitution: Resuspend in 100% ethanol to make 1.5 mg/mL stock<br>Storage: -20°C | Aliquot stock and store at -20°C for up to 6 months |
| Heparan sulfate | Stock conc: 1 mg/mL<br>Reconstitution: 1 mg/mL in $H_2O$<br>Storage: -20°C | Aliquot stock and store at -20°C for up to 6 months |
| Oleic acid | Stock conc: 0.1 mg/mL<br>Reconstitution: 0.1 mg/mL in 100% ethanol<br>Storage: -20°C | Aliquot stock and store at -20°C for up to 6 months |
| Gondoic acid | Stock conc: 0.001 mg/mL<br>Reconstitution: 0.001 mg/mL in 100% ethanol<br>Storage: -20°C | Aliquot stock and store at -20°C for up to 6 months |
| Matrigel | Use at: 1:30 | Avoid multiple freeze-thaws |
| Collagen IV | Stock conc: 200 µg/mL<br>Reconstitution: 200 µg/mL in PBS<br>Storage: -80°C | Aliquot stock and store at -20°C for up to 12 months |

**Supplementary Table 2: List of FACS antibodies**

| Antibody | Clone | Fluorochrome | Company | Catalog # | Dilution |
|---|---|---|---|---|---|
| KDR | Avas12 | PE | Biolegend | 136403 | 1:200 |
| PDGFRα | APA5 | Biolegend | Biolegend | 135911 | 1:200 |
| CXCR4 | L276F12 | Alexa Fluor 647 | Biolegend | 146504 | 1:200 |
| Kit | 2B8 | BV421 | BD Biosciences | 566290 | 1:200 |
| CD41 | MWReg30 | FITC | Biolegend | 133903 | 1:200 |
| CD16/32 | 93 | BV711 | Biolegend | 101337 | 1:200 |
| CD45 | 30-F11 | Alexa Fluor 700 | Biolegend | 103128 | 1:200 |
| CD16/CD32 (Fc Block) | 2.4G2 | Unconjugated | Bio X Cell | BE0307 | 1:50 |
| $CX_3CR1$ | SA011F11 | BV785 | Biolegend | 149029 | 1:200 |
| CD11b | M1/70 | FITC | Tonbo Biosciences | 35-0112 | 1:200 |
| P2RY12 | S16007D | PE | Biolegend | 848003 | 1:200 |
| CD45.2 | 104 | Alexa Fluor 700 | Biolegend | 109822 | 1:400 |
| CD11b | M1/70 | PE | ThermoScientific | 12-0112-82 | 1:800 |
| CD11c | N418 | APC | Biolegend | 117310 | 1:800 |
| SiglecF | S17007L | BV421 | Biolegend | 155509 | 1:400 |

**Supplementary Table 3: List of IF antibodies**

| Antibody | Host/Clone | Company | Catalog # | Dilution |
|---|---|---|---|---|
| BRACHYURY / T | rabbit/ EPR18113 | Abcam | ab209665 | 1:1000 |
| BRACHYURY / T | goat/ AF2085 | R&D Systems | KQP0618021 | 1:200 |
| FOXF1 | goat/ polyclonal | R&D Systems | AF4798 | 1:400 |
| GATA6 | rabbit/ D61E4 | Cell Signaling Technology | 5851 | 1:500 |
| VE-CADHERIN | rabbit / polyclonal | ThermoScientific | 36-1900 | 1:200 |
| RUNX1 | rabbit/ polyclonal | ThermoScientific | PA5-85543 | 1:400 |
| IBA1 | goat/ polyclonal | Novus Biologicals | NB100-1028 | 1:500 |
| IBA1 | goat/ polyclonal | FUJIFILM Wako Chemicals | 019-19741 | 1:500 |
| PU.1 | rabbit/ E.388.3 | ThermoScientific | MA5-15064 | 1:200 |
| KI-67 | rat/ SolA15 | Invitrogen | 14-5698-82 | 1:500 |
