## Supplementary Experimental Procedures for "Modeling tissue-resident macrophage development from mouse pluripotent stem cells"

### Supplementary Experimental Procedures: Step-by-step protocol for the directed differentiation of mouse EpiSCs into immature TRMs and microglia-like cells

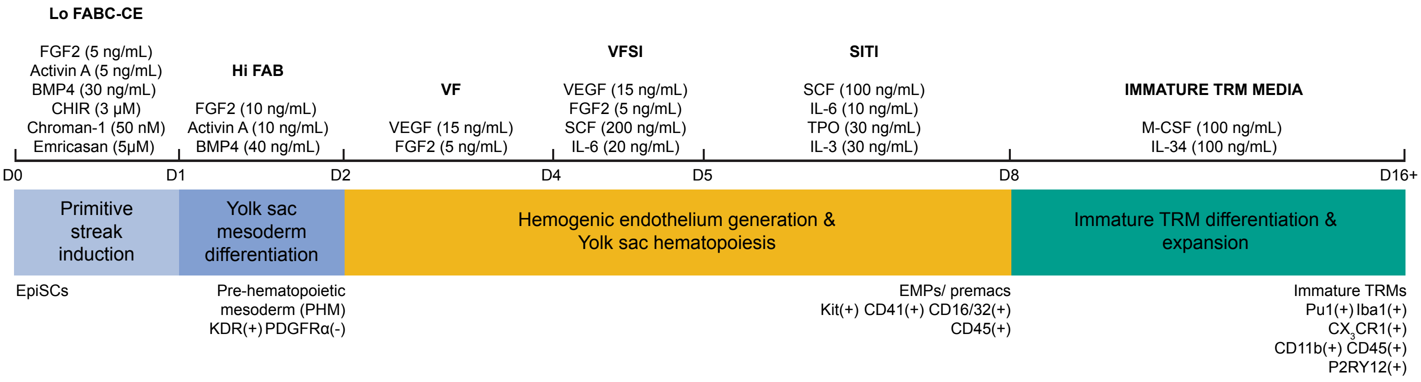


#### Notes before starting

- Prepare all media no more than two days before use.
- Before use, all media should be warmed to 37°C using a water or bead bath.
- All steps should be performed in a sterile BSC using rigorous aseptic technique.
- For conversion of mouse ESCs into EpiSCs and culture of EpiSCs, see Medina-Cano et. al., 2022.

#### Preparing fibronectin + laminin-coated cell culture dishes

1. First, coat plates with 16.7 μg/mL of fibronectin in PBS-/- (PBS without calcium or magnesium) for 30 minutes at room temperature. Ensure that entire surface of plate is covered.
2. Aspirate fibronectin and wash wells **twice** with PBS-/-.
3. Coat fibronectin-coated wells with 10 μg/mL laminin-521 in PBS+/+ (PBS with calcium and magnesium). For same day use, put laminin-filled plates in incubator at 37 °C for 2 hours or seal plates with parafilm and store at 4 °C overnight. Plates can be stored for up to one week.

#### Preparing and plating EpiSCs

1. Aspirate EpiSC media from a plate of confluent EpiSCs.
2. Wash each well twice with PBS-/-
3. Remove PBS from the plate and add 0.1 U/μL pre-warmed Collagenase IV (diluted in HBSS+/+) to each well to preferentially dissociate EpiSC colonies without feeder contamination.
    a. *Optional: place the plate with collagenase in the incubator to increase the speed of dissociation.*
    b. *Gently tap the sides of the tissue culture plate to encourage colony lifting.*
    c. *Do not leave collagenase on wells for over 25-30 minutes as this will cause feeder cells to dissociate.*
4. Once EpiSC colonies are lifted, gently collect colonies using serological pipette and transfer to a 15 mL centrifuge tube.
    a. *Optional: wash each well with HBSS+/+ to collect any remaining colonies and combine with cell solution in the 15 mL tube.*
5. Centrifuge cells at 125 g for 4 minutes to pellet EpiSC colonies and aspirate supernatant.
6. Resuspend the pellet with 1 mL Accutase to break colonies into a single cell suspension. Allow for 1-2 minutes of dissociation.
7. Once cells are completely dissociated, add 5-10 mL PBS-/- to dilute Accutase.
8. Centrifuge the solution at 200 g for 3 minutes to pellet single cells.
9. Aspirate the supernatant and resuspend in **Lo FABC-CE** media (See Supplementary Table 1).
10. Calculate the appropriate number of **live** cells to seed the desired plate, then prepare a master solution with 60,000 EpiSCs/cm$^2$. (eg. For 96-well plate, use 200ul media + ~20,000 cells per well. Scale up cell number **and** media volume of media per cm$^2$).
11. Remove the fibronectin + laminin-coated plate from the incubator and aspirate laminin solution. Do not wash plate and do not allow plate to dry – the cell suspension must be added immediately.
12. Homogenize the cell solution and immediately plate cells directly onto fibronectin + laminin-coated plates at the appropriate concentration and in the appropriate volume.

13. Place the plate in the incubator for 24 hours and then change media.

**Changing Media from D1-D8**

1. Slowly aspirate all media from all wells at a 45-degree angle to the BSC surface. Do not wash cells between media changes.
2. Add the appropriate volume of the appropriate media by pipetting slowly along the side of the well to ensure minimal cell loss (see Supplementary Table 1 for all media compositions).

**Preparing Matrigel-coated cell culture dishes**

1. Thaw Matrigel overnight on ice at 4°C. *Do this before D8 of differentiation.*
2. The next day, dilute Matrigel at a ratio of 1:30 in **cold** DMEM/F12 and add appropriate volumes of Matrigel per well.
3. For same day use, put plates at 4°C for 2 hours. Matrigel-coated plates may also be stored for up to a week at 4°C. Seal plates with parafilm to minimize evaporation.

**Splitting and replating cells at day 8**

1. Since there will be some live cells in suspension, collect all media into 15ml – 50ml centrifuge tubes. Spin down at 300g for 4 minutes to pellet non-adherent cells.
2. Then, without washing adherent cells cells, add appropriate volume of Accutase to each well and leave cells to lift and dissociate for about 1 minute.
3. Using a pipette, collect all cells from the plate and combine with pelleted non-adherent cells from step 1. Add PBS-/- to dilute Accutase (add PBS-/- at at least a 1:1 ratio of PBS:Accutase)
4. Centrifuge cells at 300 g for 4 minutes to pellet day 8 cells. Aspirate supernatant.
5. Resuspend the pellet in immature TRM media (See Supplementary Table 1) and count cells in Trypan blue.
   **NB**: Wells are highly confluent at this stage and viability is usually around 50%. Smaller wells might result in lower viability.
6. Using live cell counts, prepare a master solution containing 90,000 – 120,000 cells/cm$^2$ in immature TRM media.
7. Remove Matrigel-coated plates from 4°C, aspirate Matrigel from wells, and plate cells directly onto the Matrigel-coated plates without washing the plates.
8. Change immature TRM media every other day or 3 times a week.

**Changing immature TRM media & passaging immature TRMs**

1. To change media, use a pipette to aspirate half of the volume of media that was originally added to the well.
2. Add the same volume of media that was removed plus ~10% more media to account for evaporation.
3. Upon confluency (usually around day 16), use a micropipette to lift immature TRMs by mechanically pipetting existing media in the wells up and down until majority of the round cells are in solution. *This dissociates macrophages from remaining endothelial cells that are more firmly attached to the plate.*
4. Spin down immature TRMs at 300g for 4 minutes and aspirate supernatant.
5. Resuspend cells in fresh immature TRM media and replate onto Matrigel-coated plates at 60,000 cells/cm$^2$. Immature TRMs proliferate faster when replated at higher density; therefore, replating densities may be increased up to 120,000 cells/cm$^2$.
   *Alternatively, immature TRMs may be replated onto collagen IV-coated plates. To coat plates, add 2ug/mL collagen-IV in PBS-/- for 30 minutes at room temperature. Remove collagen-IV and plate immature TRMs at the appropriate density directly onto collagen-coated wells.*

#### Differentiating immature TRMs into microglia-like cells

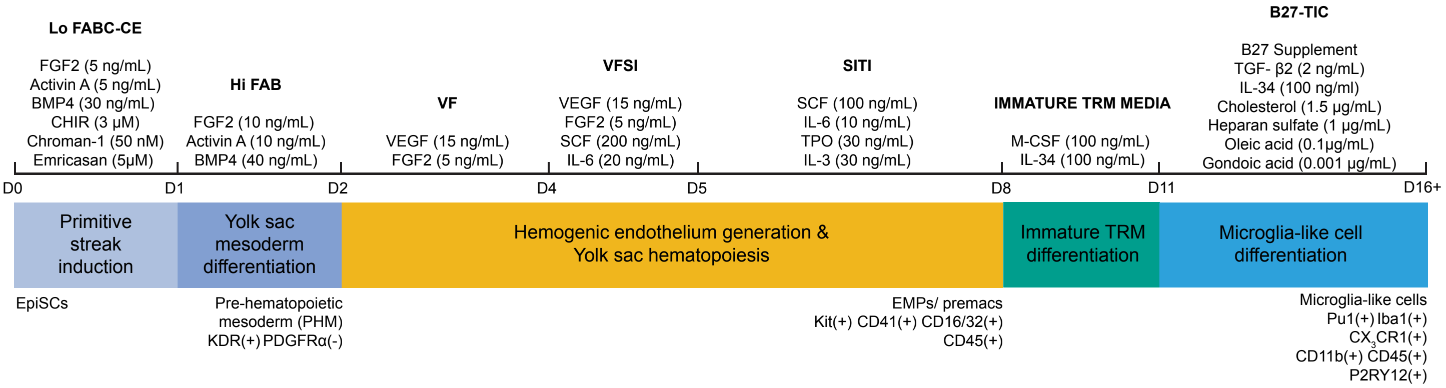


1. Follow the same steps described in the immature TRM differentiation protocol until day 11, when some immature TRM cells have begun to show up.
2. At day 11, aspirate all immature TRM differentiation media and replace with B27-TIC media.
3. Change media and passage cells in the same way that it is done for immature TRMs.

#### Potential questions and concerns

<u>Low cell density after plating</u>

- Ensure that EpiSCs were at least 70-80% confluent before beginning differentiation to guarantee more accurate cell counts.
- Ensure fibronectin, laminin and Matrigel are diluted to the right final concentration and left to coat the plates for the specified amount of time. Do not rinse culture plates after laminin or matrigel coating.
- Some cell lines have poor attachment. If this is a recurring issue for specific lines, increase the starting density when plating.

<u>Cell loss during media changes</u>

- Use one hand to tilt the tissue culture plate towards you at a 45° angle and maintain this angle while gently removing and dispensing media along the side of the well (slow ejection speed on the pipettor). This will ensure minimal loss of cells.
- Increase the size of the wells. Cell loss is more common on smaller sized wells.

<u>Low efficiency of differentiation</u>

- This protocol was optimized using EpiSCs passaged no more than 12 times. Higher passage lines may respond differently to the differentiation protocol. In general, it is best to avoid prolonged culture of EpiSCs.
- Some cell lines may require extended periods in immature macrophage media to expand and reach confluency. For such cell lines, replate day 8 cells at a higher density and/or allow for 3-4 additional days of culture.
